# Comparative epigenetic and genetic spatial structure in Mediterranean mountain plants: a multispecific comparison

**DOI:** 10.1101/2023.09.02.556052

**Authors:** Javier Valverde, Mónica Medrano, Carlos M. Herrera, Conchita Alonso

## Abstract

Epigenetic information can be heritable, but also respond to key environmental variables in situ, endowing individuals with an additional capacity to adapt to environmental changes. Thus, it is likely that in sesile organisms such as plants, part of the spatial epigenetic variation found across individuals will reflect the environmental heterogeneity of populations. Analysing the departure of the spatial epigenetic structure from the baseline genetic variation can help in understanding the value of epigenetic regulation in species with different breath of optimal environmental conditions. We performed a multispecies study that considered seven pairs of congeneric plant species, each encompassing a narrow endemic with habitat specialization and a widespread species. In three populations per species we used AFLP and methylation-sensitive AFLP markers to characterise the spatial genetic and epigenetic structures. In contrast to widespread species, narrow endemics showed a significant and generalised lower epigenetic than genetic differentiation across species. Within most populations of narrow species, epigenetic variation was less spatially structured than the genetic variation. This pattern resulted from a lack of correlation between epigenetic and genetic information in populations of narrow endemics. We argue that the differences found between narrow endemics and widespread congeners reflect contrasting breaths of environmental requirements. We pose the hypothesis that in species with a narrow niche breath, epigenetic variation may be more similar across populations and among individuals within a population given the expected higher similarity in environmental requirements.

## Introduction

In sessile organisms such as flowering plants, the distribution of genetic information in space – known as spatial genetic structure (SGS, herein)–, reflects the ubiquitous effect of gene flow determined by the most frequent short-distance dispersal of pollen and seeds. Usually, these processes result in a SGS characterised by an exponential decay of genetic relatedness with distance (Wright 1943; Epperson 1995; Rousset 2000). Characterising the SGS both among and within populations has been important in evolutionary ecology studies because of its retrospective ability to infer historical eco-evolutionary processes. For instance, contrasting SGS can reflect differences in seed dispersal (Hardy et al. 2006; Hamrick and Trapnell 2011), mating systems (Vekemans and Hardy 2004) or environmental factors such as habitat fragmentation and land use (Kloss et al. 2011; Rico and Wagner 2016; Lompo et al. 2020). The importance of characterising the SGS in wild plant populations rests on the fact that it represents the spatial template of genetic variation that defines the evolutionary potential of populations. In addition, recent studies support that plants can react to external environmental factors through epigenetic marks and alter phenotype without showing changes in their DNA sequence (e.g. Lira-Madeiros et al. 2010; Herrera and Bazaga 2011, 2016; Zhang et al. 2012; Latzel et al. 2013; Wu et al. 2013; Nicotra et al. 2015; Wilschut et al. 2016; Lehmair et al. 2022). Epigenetic changes are therefore an additional source of variability to the spatial template of plant phenotypic variation upon which selection acts (Herrera et al. 2016, 2017), and may play an important role in the fate of populations, especially in harsh environments. (Richards 2006; Jablonka and Raz 2009; Balao et al. 2018; Medrano et al. 2020). Analysing spatial epigenetic structure (SEGS) offers an opportunity to better understand the adaptive value of epigenetic variation in natural populations (Herrera et al. 2016, 2017), still there is a substantial paucity of such studies (see e.g., Lele et al. 2018; Guan et al. 2020; Shen et al. 2021).

Methylation of DNA cytosine residues is the best understood epigenetic mechanism in plants (Feng et al. 2010). Much of these epigenetic marks can be heritable for several generations (Richards 2006; Turner 2009; Verhoeven et al. 2010; Johannes and Schmitz 2019, Shi et al. 2019). Heritable epigenetic information is dispersed by the same means as genetic information (mainly seeds and pollen) and therefore the spatial distribution of both should a priori be affected in a similar way under a migration-drift equilibrium (Slatkin 1993; Herrera et al. 2016). However, in addition to genetically-determined and spontaneous methylation variants, epimutations can be directly induced by environmental changes (Gao et al. 2010; Lira-Medeiros et al. 2010; Verhoeven et al. 2010, 2016; Noshay and Springer 2021), meaning that the inherited epigenetic signature of a dispersed individual may be maintained or reset during its lifetime depending on the stability of the marker and the similarity of the offspring environmental conditions to that of its progenitors (Ibañez et al. 2021). In fact, several studies suggest that populations experiencing similar habitat conditions should express similar environmentally-induced methylation variants (Paun et al. 2010; Schulz et al. 2014).

The simultaneous study of the genetic and epigenetic spatial structures of a group of sampled individuals, as proposed by Herrera et al. (2016), can provide us with essential information about the processes that shape natural epigenetic variation in nature. Using genetic variation as a baseline null model, we can envision three main scenarios (see Figure S1). In the equilibrium between gene flow and epigenetic resetting, SGS and SEGS may show similar patterns if dispersal is the only process operating or if epigenetic reset between generations is minimal. This scenario implies a high genetic-epigenetic correlation between individuals. If, on the cotrary, environmentally driven epigenetic readjustment occurs, the SEGS will deviate from the SGS with a steeper or more gradual decline with distance depending on the grain of environmental heterogeneity, implying a decoupling of the genetic-epigenetic correlation. Taking into account the processes explained above, it is expected that with equivalent dispersal mechanisms the SEGS will depend on: 1) the range of suitable environmental conditions of each species (niche breath), and 2) the spatial distribution of the environmental variables that define such conditions (i.e., the grain of environmental heterogeneity). The use of this framework is therefore a promising tool for exploring and formulating hypotheses to help understand the interaction between species’ environmental requirements and epigenetic variation in a spatially explicit context.

Comparative studies of congeneric species pairs comprising a restricted and a widespread species provide a unique opportunity for understanding evolution of species with narrow environmental requirements (Karron 1987; Purdy et al. 1994; Jiménez-Mejías et al. 2015). In particular, narrow endemics (sensu Rabinowitz 1981) are restricted species with very specific environmental requirements that normally limit their distribution to small and isolated populations (reviewed in Slatyer et al. 2013). Theoretically, these circumstances would lead to a genetic impoverishment of their populations through drift and inbreeding (Barrett and Kohn 1991; Ellstrand and Ellam 1993; Gitzendanner and Soltis 2000). However, several studies indicate that narrow endemics from the Mediterranean depart from this prediction, showing unexpectedly high population genetic (Fernández-Mazuecos et al. 2014; Jiménez-Mejías et al. 2015; Forrest et al. 2017; Medrano et al. 2020) and epigenetic diversities (e.g. Herrera and Bazaga, 2010; Medrano et al., 2014, 2020). In addition, in contrast to species with a widespread distribution, genetic and epigenetic population diversities in Mediterranean narrow endemics have shown to be positively correlated but in such a way that in those populations with especially low genetic diversity, epigenetic diversity is unusually high (Medrano et al. 2020). These latter findings are important, as they indicate that epigenetics may act as a compensatory mechanism to create phenotypic variability. But they also suggest that the spatial correlation between genetic and epigenetic information across individuals may not behave similarly in narrow and widespread species, which may ultimately be reflected in contrasting patterns of SGS and SEGS.

Based on the shared characteristics among narrow endemics we can expect differences with widespread congeners in the SGS and SEGS. As for the smaller size of their populations, it is predicted an increased mating between nearby relatives which can be further enhanced under aggregated distribution of individuals (Doligez et al. 1998; Lara-Romero et al. 2016). Thus, following the findings in fragmented and small populations, SGS may be higher in populations of narrow endemics (e.g. Gapare and Aitken 2005; Eckstein et al. 2006; De-Lucas et al. 2009; Pandey and Rajora 2012; Rico and Wagner 2016). As for their narrower niche breath, these species inhabit highly specific habitats such as rocky outcrops that are often spatially dispersed at scales that can vary from cm to km (Lavergne et al. 2004; Thompson et al. 2005; Giménez-Benavides et al. 2018) and that condition the distribution of individuals and populations to such microenvironments (e.g. Luzuriaga et al. 2015; Schouten and Houseman 2019; Pescador et al. 2020). In contrast, the broader niche breath of widespread species allows for the colonization of diferent and more heterogeneous hábitats (Slatyer et al. 2013). Thus, if part of the epigenetic variation is environment-responsive we would expect higher independence of SEGS from the baseline SGS in narrow endemics.

Here, we explore the previous expectations in the seven pairs of congeneric plant species previously studied in Medrano et al. (2020). Each congeneric pair belongs to a different plant family and consists of a narrow endemic and a widespread species that inhabit the Sierra de Cazorla mountain range (SE Spain). This multispecies sampling scheme allows drawing taxonomically generalised conclusions about the type of species distribution (narrow endemics *versus* widespread species), while assuming other differences across species that, if relevant, will blur the potential to fulfil our expectation. Specifically, we 1) compare the SGS and SEGS across populations at a regional scale, 2) use the framework proposed by Herrera et al. (2016) to investigate the fine-scale SGS and SEGS within populations, and 3) use structured equation models and comparative analyses to test differences between endemics and widespread congeners in the spatial distance-genetic-epigenetic relationships. An understanding of the sign and intensity of the deviation of SEGS from the SGS baseline in these two groups of species will provide a better understanding of the role of epigenetics in the adaptation to harsh environments.

## Materials and methods

### Study species and field sampling

This study was carried out in the Sierras de Cazorla, Segura y Las Villas Natural Park (Jaén Province, Spain). We chose seven pairs of congeneric plant species from seven different families. Each congeneric pair consisted in a narrow endemism from the Baetic Range (restricted distribution; R, hereafter) and a widely distributed species (W, hereafter), all inhabiting this mountain range. These species correspond to those studied in Medrano *et al*., (2020): *Anthyllis ramburii* (R) and *A. vulneraria* (W); *Aquilegia pyrenaica* subsp. *cazorlensis* (R) and *A. vulgaris* (W); *Convolvulus boissieri* (R) and *C. arvensis* (W); *Erodium cazorlanum* (R) and *E. cicutarium* (W); *Daphne oleoides* (R) and *D. laureola* (W); *Teucrium rotundifolium* (R) and *T. similatum* (W); and *Viola cazorlensis* (R) and *V. odorata* (W). All species are characterised by dispersing their seeds mainly at short distances and for being pollinated by insects. Narrow endemics correspond to species with a restricted geographic distribution and inhabiting specific stressing microenvironments (Rabinowitz 1981), while widespread species inhabit a broader range of habitats (see Table S1).

For each species we chose three populations based on their accessibility and ensuring that both these and the populations of the congeneric species were located within the same geographical area (Figure 1; see Medrano et al. 2020 for exact locations). From April to June we sampled 23 to 40 widely spaced flowering individuals per population. Due to differences in population size, while sampled individuals in smaller popuations spanned most of the population spatial range, sampling in the larger populations was carried out in a representative area that was not excessively large. The chosen plants were spatially located using a GPS device (Garmin GPSMAP 64s). Spatial data for three of the populations studied that were located in canyons or under steep cliffs (two of *Aquilegia cazorlensis* and one of *Anthyllis ramburii*) were unable to be obtained, so these populations were discarded from the fine-scale spatial analyses. To guarantee GPS accuracy, two measurements were taken from the first and the last individual sampled per population. From each plant we collected 5-6 new fully expanded leaves with no sign of senescence or disease that were immediately dried in silica gel for further analyses.

**Figure 1.-.**
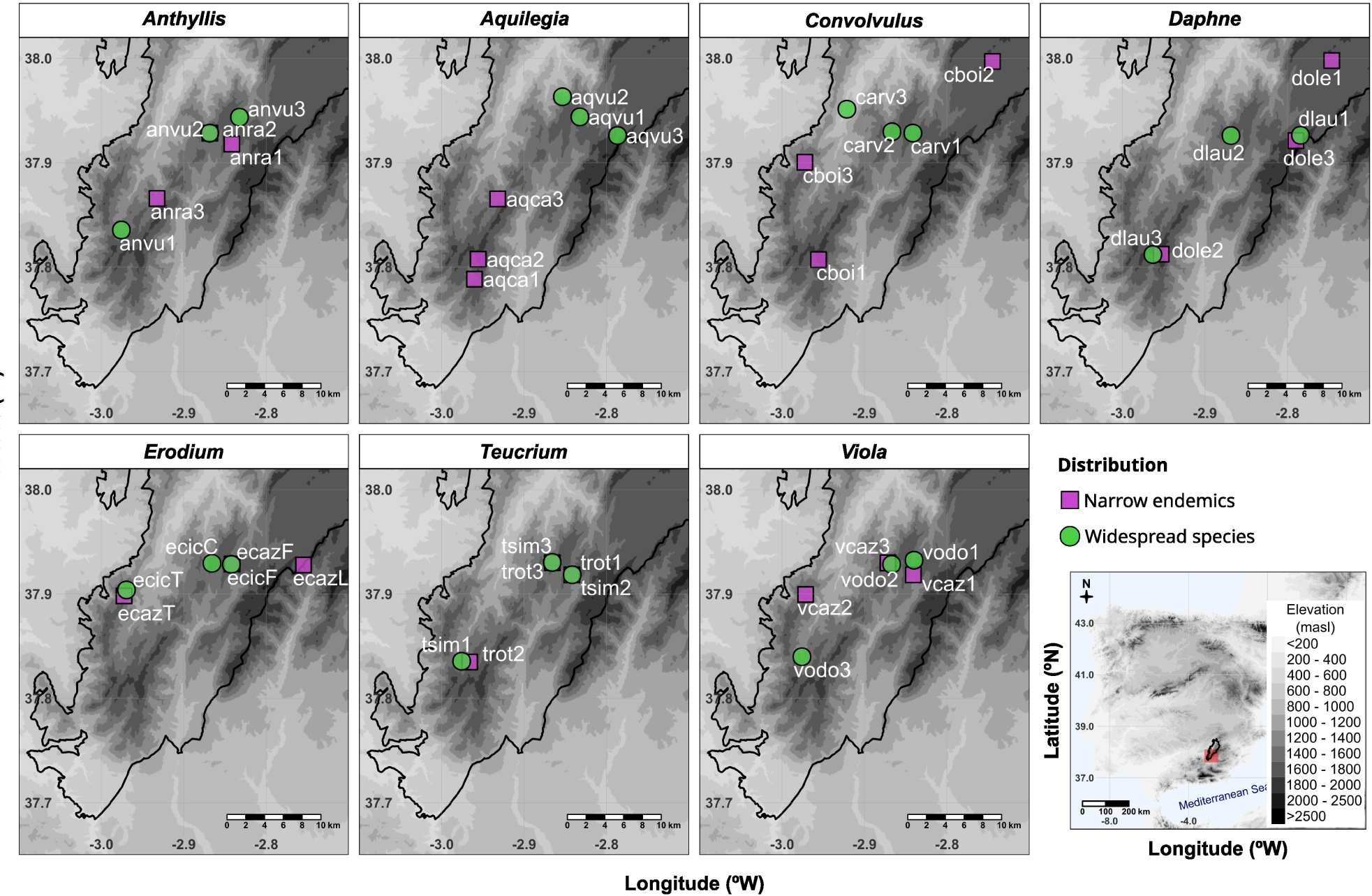
Geographical location of the studied populations at the Natural Park of Sierras de Cazorla, Segura y Las Villas in south-eastern Peninsula Iberica. For each of the studied genus the approximate location of the sampled populations of the narrow endemic and widespread congener are shown with purple squares and green circles respectively. The delimited area depicts the limits of the natural park.

### Laboratory procedures

We used amplified fragment length polymorphism (AFLP; Vos et al. 1995) and the related methylation-sensitive amplified polymorphism technique (MSAP; Reyna-López *et al*., 1997; Fulneček and Kovařík, 2014) to obtain the genetic and epigenetic fingerprint of each of the 1,088 sampled individuals. The laboratory procedures and primer combinations are detailed in Medrano et al. (2020) and in Table S2. AFLP and MSAP band scoring resulted in two binary matrices, one depicting the presence or absence of each AFLP marker and another depicting the presence or absence of each hemi- or fully methylated epiloci (Schulz *et al*., 2014). We assessed the repeatability of banding patterns for each species by repeating the entire AFLP/MSAP protocol in a number of randomly selected samples (8.6–17.6 % of samples per species for AFLP; 13.3–29.3% of samples for MSAP; see Medrano et al. 2020 for further details). We also discarded non-informative monomorphic loci that showed less than 5% of variability across samples. This yielded a variable number of polimorphic loci that varied from 86 (*Daphne oleoides*) to 418 (*Teucrium similatum*) for AFLP markers, and from 95 (*Daphne oleoides*) to 213 (*Teucrium similatum*) for MSAP markers (see Table S3 and Medrano et al. 2020 for further information).

Despite AFLP and MSAP techniques have the disadvantage of resulting in dominant bands that difficult the distinction between heterozygous or homozygous (Paun and Schönswetter, 2012), these techniques have important advantages when working with several species and large sample sizes. First, they allow reliable detection of genome-wide variants of genetic and epigenetic signatures in non-model species lacking genomic resources (Schrey *et al*., 2013; Medrano *et al*. 2014, 2020; Herrera *et al*. 2016; Wilschut *et al*. 2016; Thiebaut *et al*. 2019). In fact, the MSAP technique has been validated as an alternative to whole genome bisulphite sequencing for detecting epigenetic variation (Lauria *et al*. 2017). Second, both techniques detect anonymous markers using a similar procedure, making them mutually comparable. Finally, its relatively low cost allows for population-based studies with large sample sizes.

### Genetic and epigenetic population structure

To assess the genetic and epigenetic structure at a regional scale, we first estimated the genetic inbreeding coefficient of each population using the Markov Chain Monte Carlo procedure implemented in the *I4A* software (Chybicki *et al*. 2011). Posterior distributions were obtained using 50000 sampling steps and 10000 burning steps. This Bayesian approach takes full advantage of the power of multi-locus data and provides robust estimates of within-population inbreeding using dominant markers (Chybicki *et al*. 2011; Stone et al. 2019; García-Castaño et al. 2021). Next, for each species we calculated the genetic and epigenetic divergence between populations using Wright’s unbiased fixation index *F*_ST_ (Wright 1951) and considering the average of the previously calculated population inbreeding coefficients. These calculations were performed with the Bayesian method implemented in the *AFLP-SURV* software (Vekemans et al. 2002) using a non-uniform prior distribution. Estimated *F*_ST_ values were compared to a null distribution of values obtained after 1000 permutations of individuals across populations that depicted a complete absence of structure among populations. An additional hierarchical analysis of molecular variance (AMOVA) was performed to partition the genetic and epigenetic variance between and within populations in each plant species using the R package ade4 (Dray and Dufour 2007).

For each group of species (narrow endemics and widespread species), we compared the genetic and epigenetic *F*_ST_ values using a non-parametric Wilcoxon paired test. To be confident of our results these analyses were repeated for *F*_ST_ values that were calculated considering inbreeding estimates of 0 (panmixia) and 1 (autogamy) that encompass the maximum range of variation.

### Fine-scale spatial genetic and epigenetic structure

To study the genetic and epigenetic structure within each of the sampled populations we calculated the genetic relatedness and epigenetic similarity between each pair of individuals using the kinship coefficient proposed by Hardy (2003) for dominant markers. The calculation of this coefficient included the required estimates of inbreeding coefficient for each population. This kinship coefficient expresses the degree of similarity between individuals relative to the average similarity found in the population. Thus, positive and negative values depict respectively higher and lower relatedness than random individuals from the population. We visually explored the fine-scale genetic and epigenetic spatial structures (SGS and SEGS) by calculating the average kinship coefficients (*F*_N_) for six distance classes that showed even number of pairwise distances and that avoided spatial gaps. The significance of each *F_N_* with respect to the expected in the absence of structure was tested using 1000 permutation and two-sided tests. These calculations were performed using the SPAGeDi software (Hardy and Vekemans, 2002).

For each population we calculated the slope of the linear relationship between the logarithm of the spatial separation between each pair of individuals and their (epi)genetic kinship coefficients (*b*). Next, we used mixed-effects models to explore whether there was any generalised trend in the relationship between SGS and SEGS. These models were applied for the group of populations of endemic plants and widely distributed plants independently. The slopes of the log_10_(distance)-kinship relationship were used as dependent variable and modelled as a function of the type of marker (AFLP or MSAP), and species was included as a random factor. We followed a model selection approximation in which the previous model was compared with a reduced model without the type of marker as explanatory variable. This comparison was based on Akaike and Bayes information criteria (AIC and BIC) and analysis of variance. To control for possible effects of the disparity in the spatial distribution of sampled individuals (Doligez et al. 1998), we included the coefficient of variation and the maximum value of the pairwise spatial distances of each population as covariates (Nagamitsu et al. 2019).

### Comparison between narrow endemics and widespread species

We explored the differences between narrow endemics and widely distributed congeners in the relationship between spatial distance, genetic relatedness and epigenetic similarity. To do so, we first defined a causal model that relates the previous three variables and that includes the following direct causal effects (Figure S2): a) the effect of the log10(distance) on genetic relatedness, which typically denotes gene dispersal (Wright 1943); b) the effect of genetic relatedness on epigenetic similarity, which reflects the dependence epigenetic marks on genetic identity (Noshay and Springer 2021); and c) the effect of the log10(distance) on epigenetic kinship, which may describe the putative dependence of environmentally-biased epigenetic marks with space (Herrera et al. 2016).

Finally, this causal structure allowed us to calculate the indirect effect of spatial distance on epigenetic kinship through genetic kinship (a + b) which reflects the epigenetic reset across space. We fitted this causal model to each of the study populations separately using structural equation models using the piecewiseSEM R package (Lefcheck 2016). This approximation allowed us to calculate the previously detailed direct and indirect effects as standardized estimates and as well as their respective confidence intervals and significance after 1000 bootstrap steps using the semEff R package (Murphy 2022). As a result, for each effect we obtained a set of estimates, each from one of the studied populations.

We tested the differences between endemics and widespread congeners in the previous direct and indirect effects by means of mixed-effects linear models. Through model selection, we compared a reduced model to a full model in order to tested to what extent the addition of the type of distribution best explained each of the estimates. Genus was included as random intercept effect in both models. Additionally, we included a random slopes model in the model selection. To control for the spatial distribution of plants in each population, all models included the coefficient of variation and maximum distances as covariates. Models yielding singular fits due to low variance in the random effects were modelled using simple linear models.

## Results

### Genetic and epigenetic population structure

Species-averaged inbreeding estimates ranged between 0.04 (*Viola cazorlensis)* and 0.96 (both species of *Teucrium*). Most species showed small although significant *F*_ST_ values for genetic and epigenetic markers (*F*_ST_ < 0.27 and *F*_ST_ < 0.37 for genetic and epigenetic, respectively; Table S4). These results are in agreement with those from the AMOVA analyses, that showed significant population differentiation although most of the genetic and epigenetic variance occurred within population (range = 62.5–99.1% for AFLP and 51.1–94.3% for MSAP; Table S5). For most of the species studied, the variation attributable to differences among populations was greater for genetic than for epigenetic markers (Figure 2). However, only in narrow endemics genetic *F*_ST_ was significantly higher than epigenetic *F*_ST_ (*V* = 28, *p* = 0.016 in endemics; *V* = 21, *p* = 0.297 in widespread species). These comparisons were consistent when considering inbreeding coefficients of 0 (*V* = 28, *p* = 0.016 in endemics; *V* = 20, *p* = 0.375 in widespread) or 1 (*V* = 27, *p* = 0.031 in endemics; *V* = 21, *p* = 0.297 in widespread), thus justifying the performance of our inbreeding estimates in these analyses.

**Figure 2.-.**
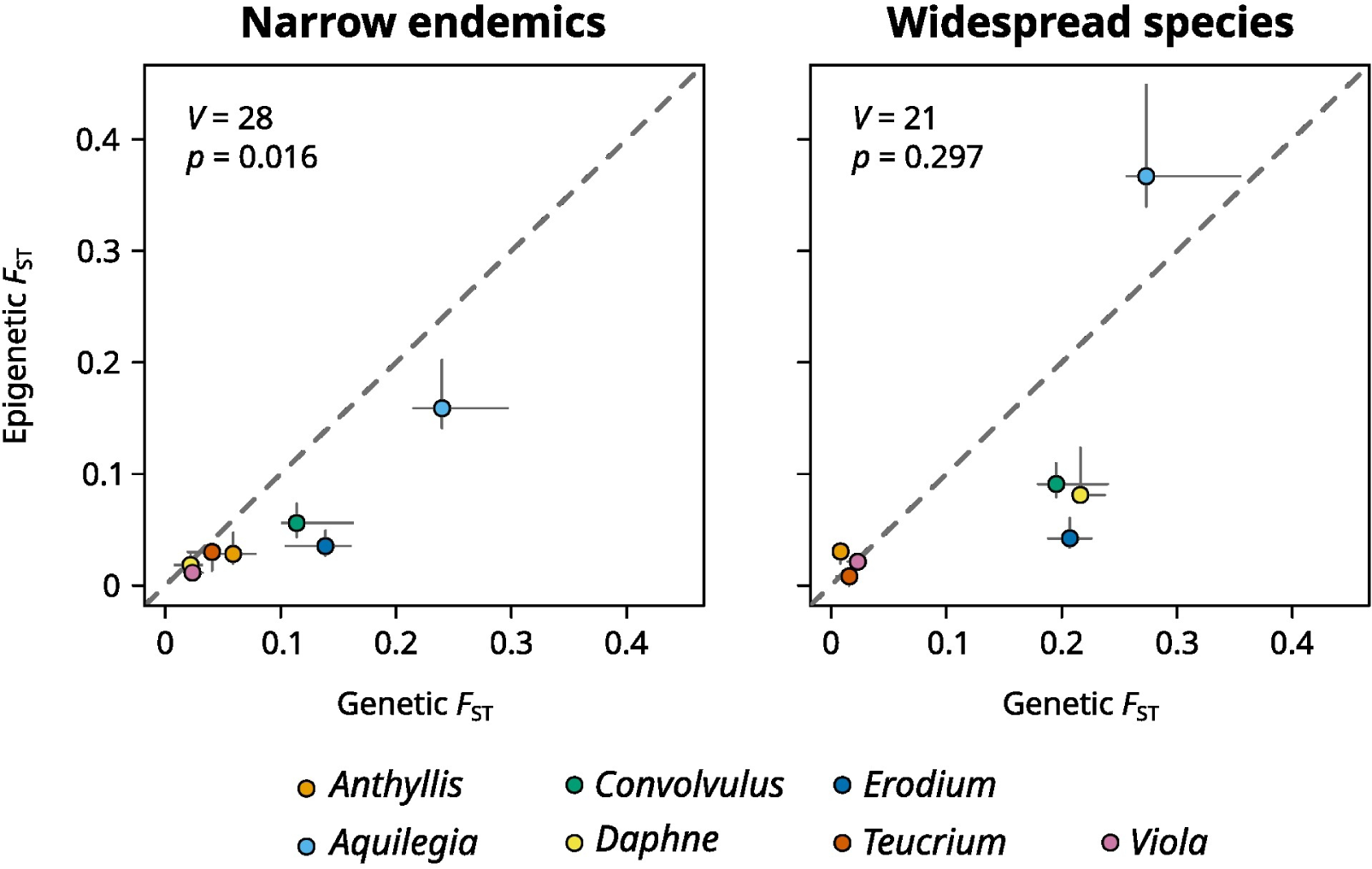
Genetic and epigenetic structure between populations. For each study species the genetic (AFLP) divergence between populations is plotted against the methylation-sensitive epigenetic (MSAP) divergence, both calculated as the Wright fixation index (FST). Error bars span FST values calculated using inbreeding coefficients of 0 and 1, while points represent values of FST calculated using the inbreeding estimates after the method proposed by Chybicki et al. (2011). The dashed line is used to contrast cases where the differentiation is relatively greater for genetic markers (lower area) or for epigenetic markers (upper area). Results from Wilcoxon paired tests for the differences between genetic and epigenetic FST values are shown in each panel.

### Fine-scale genetic and epigenetic spatial structure

The populations sampled showed a varied spatial distribution of individuals (see examples in Figure 3) that resulted in a high heterogeneity in the distribution of plant-plant spatial distances (CV range = 0.48–1.17; see Figures 3 and S3), with 8 and 5 populations of endemic and widespread species respectively showing a multimodal distribution of spatial distances (Hartigans’ multimodality test p<0.05). A total of 24 populations displayed a significant decrease of genetic relatedness with log_10_(distance) (*b* = −0.11 – −0.01, *p* ≤ 0.047), while for epigenetic markers this relationship was significant and negative in 14 populations (*b* = −0.05 – −0.01, *p* ≤ 0.034; Figure S3; Table S6).

**Figure 3.-.**
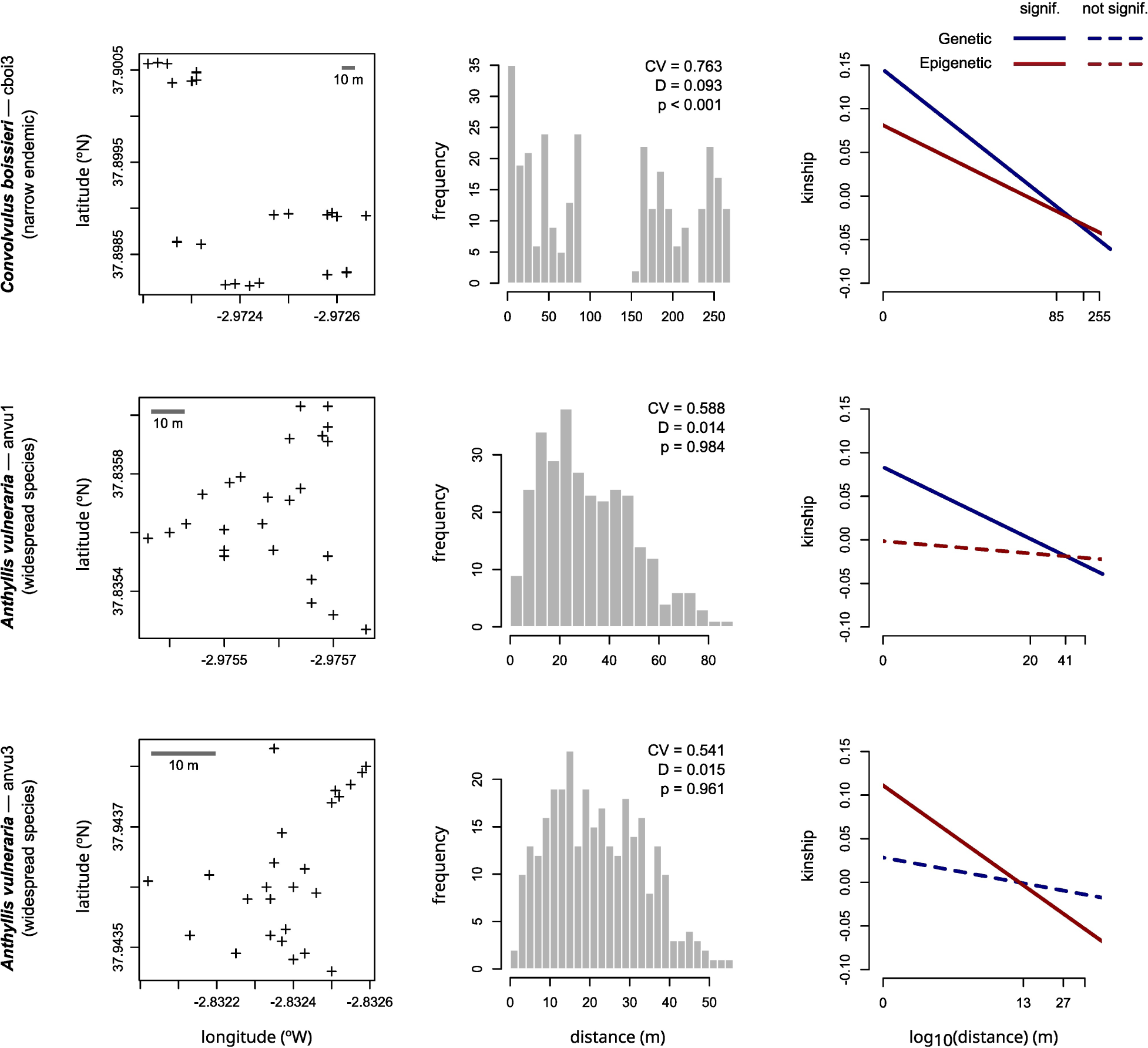
Spatial distribution of sampled individuals and spatial genetic and epigenetic structure in three of the studied populations. These populations have been arbitrarily chosen to exemplify the variability found across populations. Left panels show the spatial distribution of sampled individuals. Middle panels depict the distribution of plant-plant spatial distances and information about its variability (CV) and multimodality (Hartigan’s D and associated p value). Right panels show the slopes of the relationship between the logarithm of the spatial distance and the genetic relatedness (blue lines) and the epigenetic similarity (red lines). Figure 3S provides all this same information for all the sampled populations.

The addition of the type of marker as explanatory variable improved the prediction of the slope of the log_10_(distance)–(epi)genetic kinships in both groups of species (see results of model selection in Table S7), which indicates differences between the genetic and epigenetic spatial structures. In this sense the marginal means of the selected models showed steeper slopes for the genetic relatedness in both narrow endemics (−0.016 ± 0.003 for genetic, and −0.009 ± 0.003 for epigenetic; Figure 4; Table 1) and widespread species (−0.030 ± 0.006 for genetic, and −0.019 ± 0.006 for epigenetic), although this comparison was not significant in widespread species.

**Figure 4.-.**
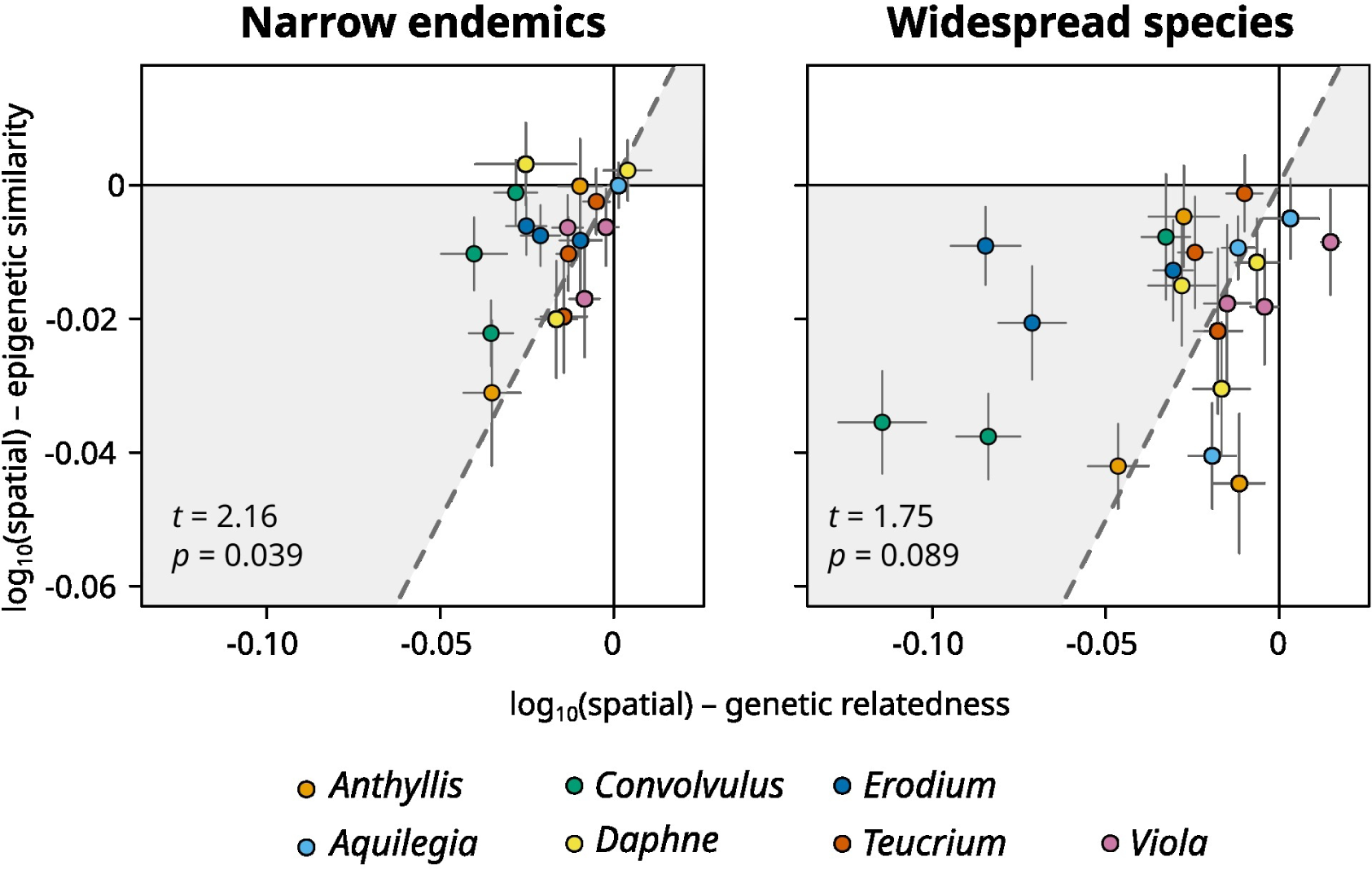
Fine-scale genetic and epigenetic spatial structure in populations of narrow endemics and widespread species. The panels summarise the slopes of the log10(distance)-genetic relatedness and log10(distance)-epigenetic similarity relationships in all populations studied. Points within the grey areas denote populations where epigenetic similarity decreases with distance less steeply than genetic relatedness (and the opposite for white areas). Error bars span ±1 SE after a jackknifing procedure over loci. Statistics at each panel refer to the term defining the type of marker in the mixed model and thus reflect overall differences between genetic and epigenetic slopes.

**Table 1.-.**
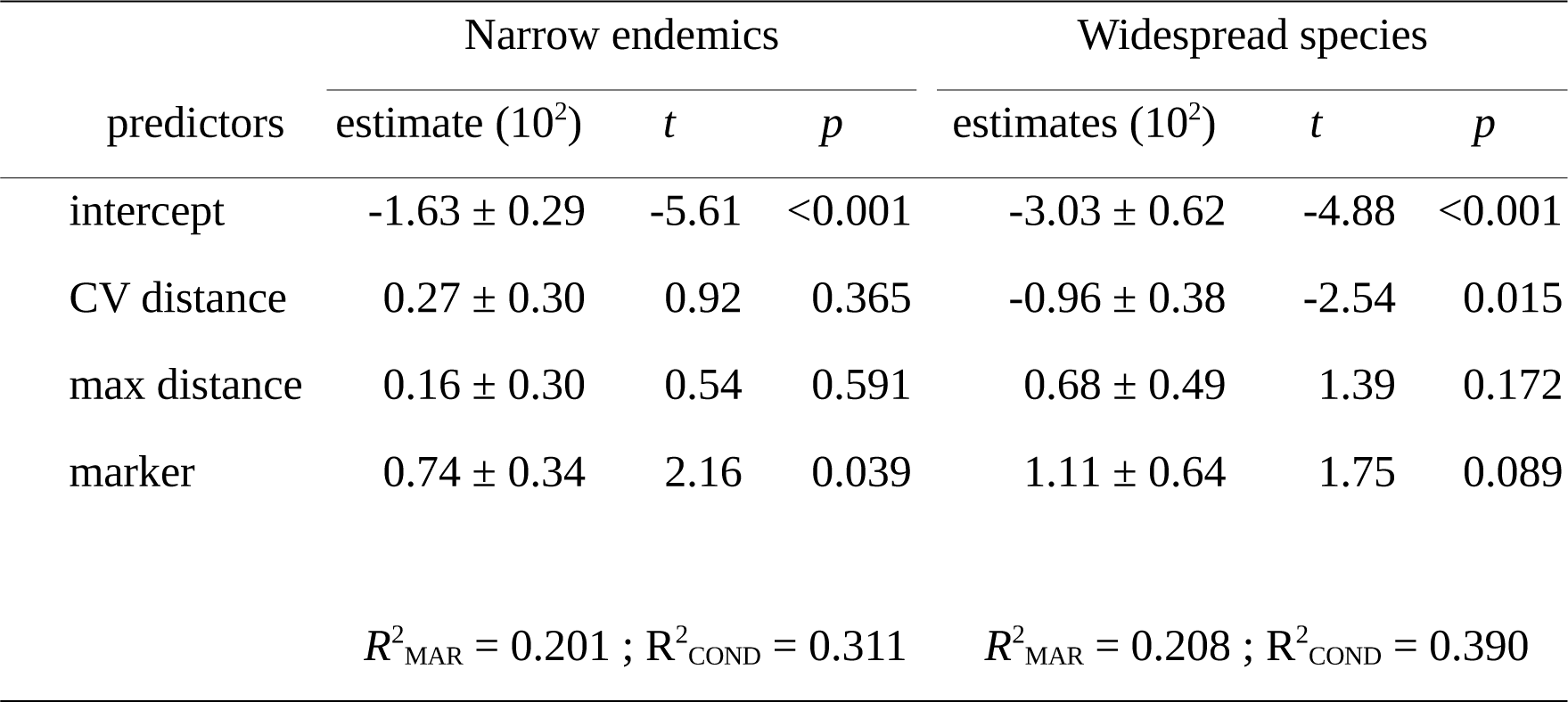
Comparison of the genetic and epigenetic fine-scale spatial structure in the studied populations from the narrow endemics and the widespread species. Models show the effect of the type of marker (AFLP or MSAP) on the slope of the relationship between logarithm of spatial distance and the (epi)gentic kinship. These random intercept models (see table S6 for model selection) include the coefficient of variation and maximum value of distance between pairs of plants in each population as covariates. Goodness-of-fit of each model is shown by the marginal and conditional R-squared statistics.

### Comparison between narrow endemics and widespread species

Structural equation models showed that the effect of genetic relatedness on epigenetic similarity was positive and significant in 55% of the populations, most of them from widespread species (4 in endemics, 12 in widespread). It also revealed that the direct effect of spatial distance on epigenetic kinship was only significant in nine out of the 39 studied populations (Table S8), although this relationship was negative in most cases. In contrast, the indirect effect of spatial distance on epigenetic kinship –mediated through genetic kinship–, was negative and significant in 41% of populations, most of them from widespread species (4 in endemics, 12 in widespread).

After model selection, the models that included the type of distribution as explanatory variable were those that best explained the direct effect of genetic relatedness on epigenetic similarity as well as the indirect effect of spatial distance on epigenetic similarity (Table S9). The estimated marginal means showed a four-times higher and significant effect of genetic relatedness on epigenetic similarity in widespread species that in their endemic congeners (0.094 ± 0.037 in endemics; 0.347 ± 0.034 in widespread, Figure 5). The indirect effect of spatial distance on epigenetic similarity was nearly eight times stronger in widespread species (−0.009 ± 0.012 in endemics; −0.068 ± 0.026 in widespread, Figure 5). In contrasts to this, reduced models were the best in explaining the direct effect of spatial distance on both genetic relatedness and epigenetic similarity (Table S9), which resulted in the same predicted means for restricted and widespread species (−0.175 ± 0.005 for the log_10_(distance)-genetic kinship and −0.055 ± 0.010 for the log_10_(distance)-epigenetic kinship, Figure 5).

**Figure 5.-.**
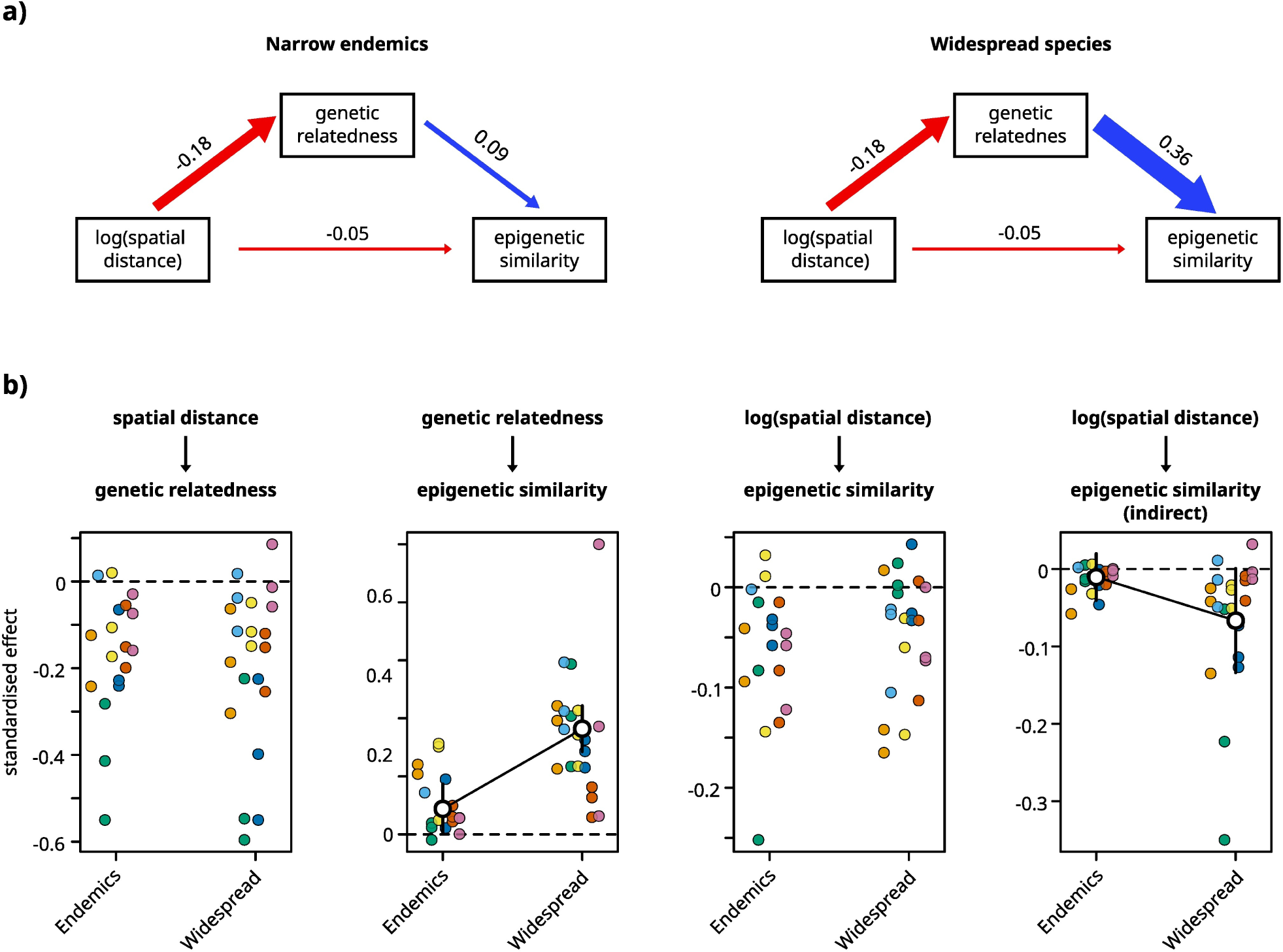
Comparison between restricted and widespread species in the relationship between spatial distance, genetic relatedness and epigenetic similarity. a) Causal model for the restricted and widespread species. The arrows show the direct causal relationships. Their size is relative to the estimated effect across populations, which is also shown. Blue and red arrows denote positive and negative values respectively. b) Comparison of the direct and indirect effects. Each panel shows the distribution of the estimated effects in populations from restricted and widespread species. Each genus is represented by a colour (please refer to the legend in Figures 2 and 4). In those effects where significant differences were found between narrow endemics and widespread species, marginal means and associated standard error bars are shown.

## Discussion

This research illustrates the broad variety of spatial structuring of genetic and epigenetic variation in natural populations of Mediterranean montane areas. Using a multispecies comparative approach with a set of seven pairs of congeneric species differing in their range of geographic distribution and habitat specialization, we tested the specific hypothesis that in species with greater habitat specificity spatial epigenetic structure should be more independent of spatial genetic structure. Our results showed that, in general, epigenetic spatial structure was lower than genetic structure both between and within populations of narrow endemisms. Within populations, such differences appear to be the result of a greater independence of epigenetic and genetic variations. To our knowledge this is the first study in comparing the spatial genetic and epigenetic structures in a group of congeneric species with contrasting distribution and using a spatially explicit approach. In the following paragraphs we discuss our findings among and within populations, and propose a hypothesis for future studies based on the patterns found at the fine scale.

### Genetic and epigenetic population structure

Our sampling scheme showed low genetic differentiation among populations in the study species, with most of the genetic variance found within populations. Wright fixation index for divergence of the studied populations at intraspecific level ranged between 0.02 and 0.24 in narrow endemics and between 0.01 and 0.27 in widespread species. These values lay within the lower part of the range reported for 86 insect-pollinated temperate plant species (average = 0.21; 95% CI = 0.01 – 0.51; Gamba and Muchhala 2020 and references therein). Caution should be taken when interpreting these parameters because an analysis of three populations per species might not be representative of population differentiation in the full geographic distribution range (e.g., Castilla et al. 2012 for *Daphne laureola*; Daco et al. 2022 for *A. vulneraria*; but see Herrera and Bazaga 2008 for *Viola cazorlensis*). However, the *F*_ST_ values obtained through the multispecies framework and population level approach may provide representative data for the particular geographic area in which the sampling was conducted. Specifically, it suggests that in our study area, the magnitude of population differentiation of endemic and widespread species could be similar, although future studies with more species and populations will be required to support these results.

Epigenetic differentiation was also low (*F*_ST_ range = 0.012–0.159), but more importantly, it was lower than genetic differentiation in most of the species studied. Especially in the endemic species, epigenetic differentiation was significantly lower than genetic differentiation, while widely distributed species did not show a significant trend across species and populations. To date, the few research on epigenetic differentiation of populations point to a lack of common pattern in the comparison between epigenetic and genetic differentiation (e.g., Wu et al. 2013; Herrera et al. 2017 *versus* Avramidou et al. 2015; Lele et al. 2018). However, in most studies where there is higher epigenetic than genetic differentiation among populations, the former is associated with environmental factors. Among these studies, it is worth highlighting those that show an association with population soil variables such as moisture, pH or chemical composition (Lira-Medeiros et al. 2010; Shen et al. 2021; Lehmair et al. 2022), which together with experimental studies controlling for soil variables (reviewed in Alonso et al. 2019), add to the evidence of the ubiquitous effect of soil characteristics and habitat features on epigenetic variation (see also Paun et al. 2010, Schulz et al. 2014). In our study, endemic species are characterised by having a narrow niche breath which is sparsely distributed in space and that determines the patchy distribution of these species. We believe that the similar habitat conditions that individuals of each species inhabit may lead to a convergence of the methylation patterns, which may explain the significantly lower epigenetic differentiation among populations.

### Fine-scale genetic structure within populations

Within most of the populations studied, the genetic variation was spatially structured, showing a significant decrease with spatial distance. This fine-scale SGS is expected for sessile organisms with short-distance dispersal mechanisms (Vekemans and Hardy 2004) such as those shared among the studied species. Narrow endemics and widespread species did not show global differences in the fine-scale SGS. This finding is somehow surprising given that narrow endemics are comprised of smaller and isolated populations. In small populations with spatially restricted pollen flow and seed dispersal, kinship between close individuals is expected to increase and, consequently, the strength of fine-scale SGS (Heywood, 1991; Vekemans and Hardy, 2004; Gapare and Aitken 2005; Volis et al. 2016). A plausible explanation of the lack of differentiation in SGS could lie in the different pollinators visiting the species studied. For instance, the endemic *Viola cazorlensis* is mainly pollinated by the hawkmoth *Macroglossum stellatarum* which is able to fly long distances and thus create a flow of pollen between individuals located far apart within the same population. On top of this, Valverde et al. (unpublished) have found a covariation between pollinator diversity and population genetic diversity in a subset of these same endemic species. It is likely that in these endemic species, the increase in foraging types that comes with high pollinator diversities promotes greater pollen movement, which in the long run may relax the SGS. This hypothesis and other related to factors conditioning reproductive patterns should further be explored, as they are key in generating genetic spatial variation (Ennos 2001; Vekemans and Hardy 2004; Gamba and Muchhala 2022).

### Patterns of spatial-genetic-epigenetic relationships

Epigenetic similarity within populations decreased with distance more gradually than genetic relatedness in most cases (i.e., steeper slope for genetic markers), indicating that epigenetic variation was in general less spatially structured than genetic variation (SEGS < SGS). Despite this trend, there were exceptions in our data, mainly in populations of some widely distributed species. For instance, opposite to the general trend, in all sampled populations of *Viola odorata* epigenetic variation was more spatially structured in the three populations. In *Anthyllis vulneraria* and *Daphne oleoides*, there was a lack of consistency in the spatial epigenetic patterns across populations. Especially, the relationship between log_10_(distance) and epigenetic similarity varied more in the widespread species, which may be due to multiple factors, ranging from environmental heterogeneity to different epigenetic responses, either independent of the environment, highly dependent on genetic variation or highly dependent on the environment.

In contrast, populations from narrow endemics showed a higher consistency across populations in the previously explained deviations of the epigenetic from the baseline genetic spatial patterns. According to the framework of Herrera et al. (2016), this suggests a more prevalent epigenetic reset unresponsive to environmental variation in these species. This reasoning is based on the fact that environmental variables usually exhibit a strong spatial autocorrelation (i.e. a decrease in environmental similarity with distance; Legendre 1993). Under such an assumption the framework predicts that while a higher epigenetic structure could be given by a spatial correlation with a spatially structured environmental variable, a lower epigenetic structure -which is our case-, would be the consequence of an epigenetic readjustment independent of the environment. However, we believe that this argument does not fit our particular case, and that, similar to what we have previously discussed about the epigenetic differentiation between populations, within-population epigenetic variation in endemic species could be explained by their habitat specificity and the expected lack of spatial structure in the microenvironmental growing conditions, as we expose below.

In the studied endemic species we found a low or even a lack of dependence of epigenetic variation on genetic variation, which contrasts with the highly significant genetic-epigenetic correlation found in widespread species. Despite that studies in the model species *Arabidopsis thaliana* demonstrate the interdependence between genetic and epigenetic signatures (Zilberman et al. 2007; Dubin et al. 2015; Kawakatsu et al. 2016), other studies demonstrate a lack of such interdependence in natural populations of non-model species (e.g. Herrera and Bazaga 2010; Richards et al. 2012; Schulz et al. 2014; Foust et al. 2016). Such contrasting findings may be due to potential higher rates of epigenetic spontaneous mutations in certain species in comparison to that of *A. thaliana*. However, evidence of the existence of epimutations associated with the environment (e.g. Lira-Madeiros et al. 2010; Herrera and Bazaga 2011, 2016; Nicotra et al. 2015; Gáspár et al. 2018) suggest that environmental stochasticity may be a major determinant of epigenetic variation. Our study is the first in identifying an association between the variation in the genetic-epigenetic correlation and a plant ecological characteristic. It is likely that in the group of endemic plants the decoupling of epigenetic variation from the baseline genetic variation is a result of an epigenetic readjustment to accommodate plant individuals to the narrow environmental conditions in which they live. However, the contrast with the alternative hypothesis of a higher rate of spontaneous epimutation associated to harsh environments (Johannes and Smith 2019) merits further study.

On top of this, our causal models corroborated the differences in fine-scale epigenetic spatial structure between endemics and widespread congeners. But more importantly, it demonstrated significant differences between endemic and widespread populations in the indirect effect of spatial distance on epigenetic similarity when mediated by the genetic relatedness of individuals. Specifically, the results indicate that in endemics the low epigenetic spatial structure within populations is mainly due to the aforementioned lack of relationship between epigenetic and genetic variation at this scale. This finding contrasts with that in widespread species, where the greater epigenetic spatial structure is mediated by the close dependence of epigenetic variation on genetic relatedness. As a result, both groups of plants differed in the fine-scale SEGS deviation from the baseline SGS.

This further supports our rationale that the environmental requirements defining each group of studied species shape the epigenetic response in space: On the one hand, the endemic plants studied inhabit highly specific microsites, mainly dolomitic rocky outcrops surrounded by a matrix of less favourable environmental conditions (e.g. soil conditions or increased interspecific competition), which is reflected in the aggregate spatial distribution of individuals. On the other hand, widely distributed species may inhabit a wider range of microsites, occupying a large fraction of space despite environmental heterogeneity. As a consequence of the higher specificity in endemic species, the environmental similarity between any pair of plants is expected to be higher than in widely distributed species. Assuming that epigenetic variation can respond to environment, epigenetic changes in endemics will be minimal, implying a homogenisation of the epigenetic similarity between individuals plants that would result in the observed decoupling between spatial epigenetic and genetic structures.

Certainly, to be fully confident of this causality, further studies should consider to characterise the microenvironmental conditions of individual plants and to incorporate this as a direct effect on epigenetic similarity in the causality model as it has been previously done in studies at the regional level (e.g. Herrera and Bazaga 2011; Lele et al. 2018). This piece of information is especially relevant given the effect that environmental distance can have on the spatial distribution of epigenetic variants (Herrera et al. 2017). In any case, our findings are in line with previous findings in which a decoupling of the genetic and epigenetic variation are accompanied by a correlation of epigenetic variation with environmental conditions (Lehmair et al. 2022) and highlight the importance of such in determining the spatial template of genetic and epigenetic variation.

## Conclusions

The study of epigenetic variation in natural populations is a promising tool for proposing and answering hypotheses of profound evolutionary significance. The diversity provided by part of epigenetic regulation can facilitate individuals to mould to environmental changes. This capacity certainly contributes to the species’ niche breadth determined by genetic variation. In our work, we have found epigenetic variation to be lower then genetic variation among and within populations of species with very specific environmental requirements and small populations. Within populations, the decoupling of epigenetic and genetic variation in these species resulted in an erosion of spatial epigenetic variation. Given the narrow environmental conditions of these species, we hypothesize that the lower spatial epigenetic structure found may reflect a spatial homogenization of environment-related epigenetic marks.

## Acknowledgements

We are very grateful to Pilar Bazaga and Esmeralda López-Perea for laboratory assistance. Noelia Zara for her help in sampling *Erodium* individuals. We thank the Consejería de Medio Ambiente, Junta de Andalucía, for authorizing this research. This work was supported by the Consejería de Innovación Ciencia y Empresa, Junta de Andalucía [P18-FR-4413] and the Ministerio de Ciencia e Innovación, Spanish Government [PID2019-104365GB-I00/AEI/10.13039/501100011033 and SUMHAL-LIFEWATCH-2019-09-CSIC-13, through European Regional Development Funds POPE 2014-2020].

## Author contributions

CA, CMH and MM designed the methodology and collected the data; CA, CMH, MM and JV elaborated the hypotheses; JV analysed the data and led the writing of the manuscript. All authors contributed to the revision of the manuscript and gave their final approval for publication.

## Data archiving

Genotype data will be made available on acceptance.

## Notes

### Competing Interest Statement

The authors have declared no competing interest.

