## Supplementary Figures for "Comparative epigenetic and genetic spatial structure in Mediterranean mountain plants: a multispecific comparison"

**Figure S1.-** Different scenarios of spatial genetic and epigenetic structures. Following the framework of Herrera et al. (2016), under a base-line inverse relationship between genetic relatedness and distance, the following relative relationships between epigenetic similarity and spatial distance can be found: a) Common slopes in the genetic and epigenetic spatial structures when dispersal is the only process operating or when epigenetic reset between generations is minimal; b) More moderate slope in the epigenetic spatial structure, or c) steeper slope in the epigenetic spatial structure whenever there is an environment-driven epigenetic reset. The deviation of SEGS from the baseline SGS should depend on the spatial grain of environmental heterogeneity.

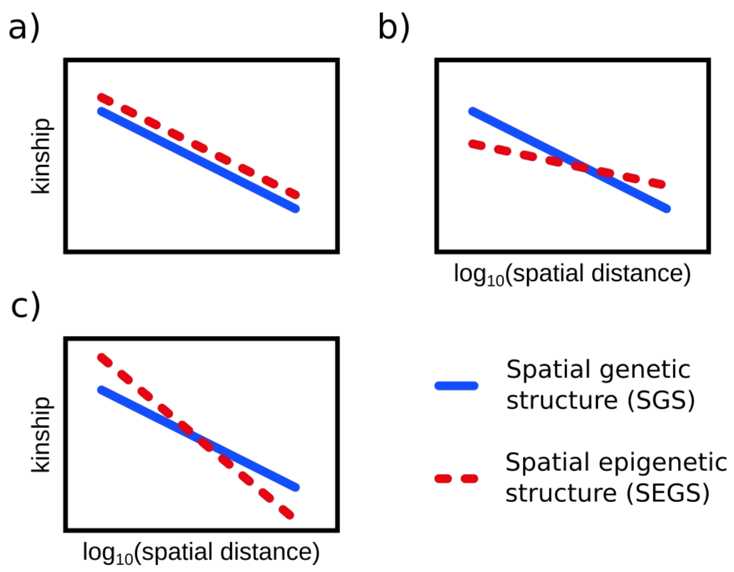

**Figure S2.-** Causal model depicting the relationships between spatial distance, genetic and epigenetic kinship. Direct causal effects: a) the effect of spatial distance on genetic relatedness, which denotes gene dispersal; b) the effect of genetic relatedness on epigenetic similarity, which reflects the dependence epigenetic marks on genetic identity; c) the effect of spatial distance on epigenetic kinship, which may describe the putative dependence of environmentally-biased epigenetic marks with space. Indirect effect of spatial distance on epigenetic kinship through genetic kinship (a + b) which reflects the epigenetic reset across space.

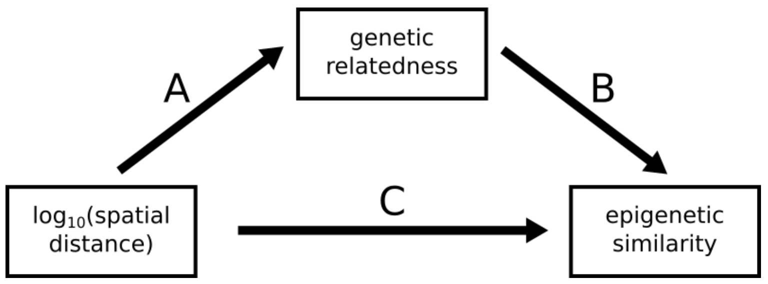

**Figure S3 (1/10).-** Spatial distribution of sampled individuals and spatial genetic and epigenetic structure within study populations. From left to right: 1) Spatial distribution of sampled individuals. 2) Distribution of distances between individuals and information on its variability (CV) and multimodality (Hartigan's D and associated p-value). 3) Mean genetic and epigenetic kinship in each distance class. Filled symbols indicate values significantly higher ($p < 0.01$) than expected after random permutation. 4) Genetic and epigenetic spatial structures. Symbols represent the average kinship between individuals from the first distance class ($F_1$) and the associated error bars represent ±1 SE after a jackknifing procedure on the loci. $\log_{10}$(spatial distance)-(epi)genetic slopes are shown with a solid line if the relationship is significant.

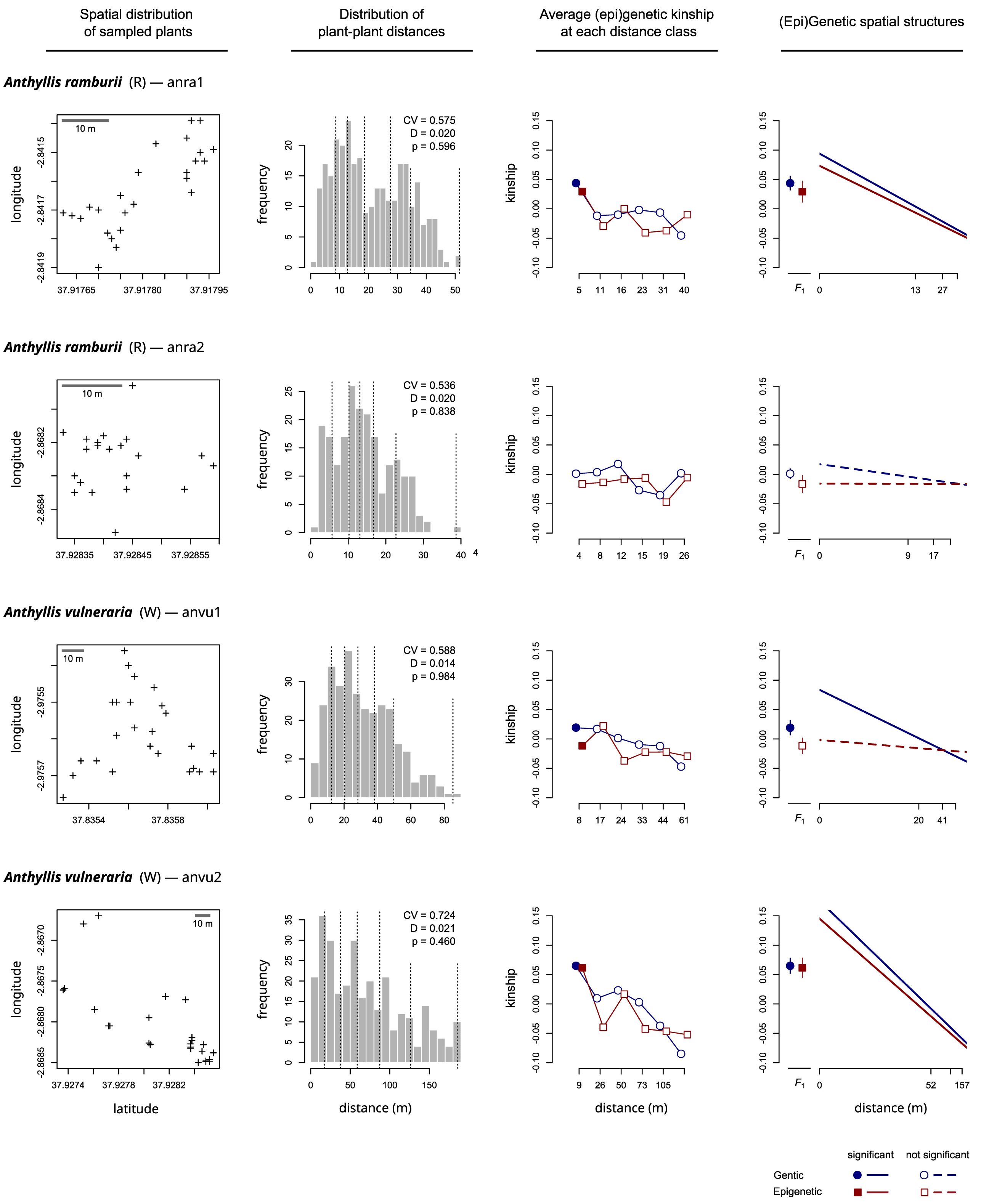

**Figure S3 (2/10).-** Spatial distribution of sampled individuals and spatial genetic and epigenetic structure within study populations.

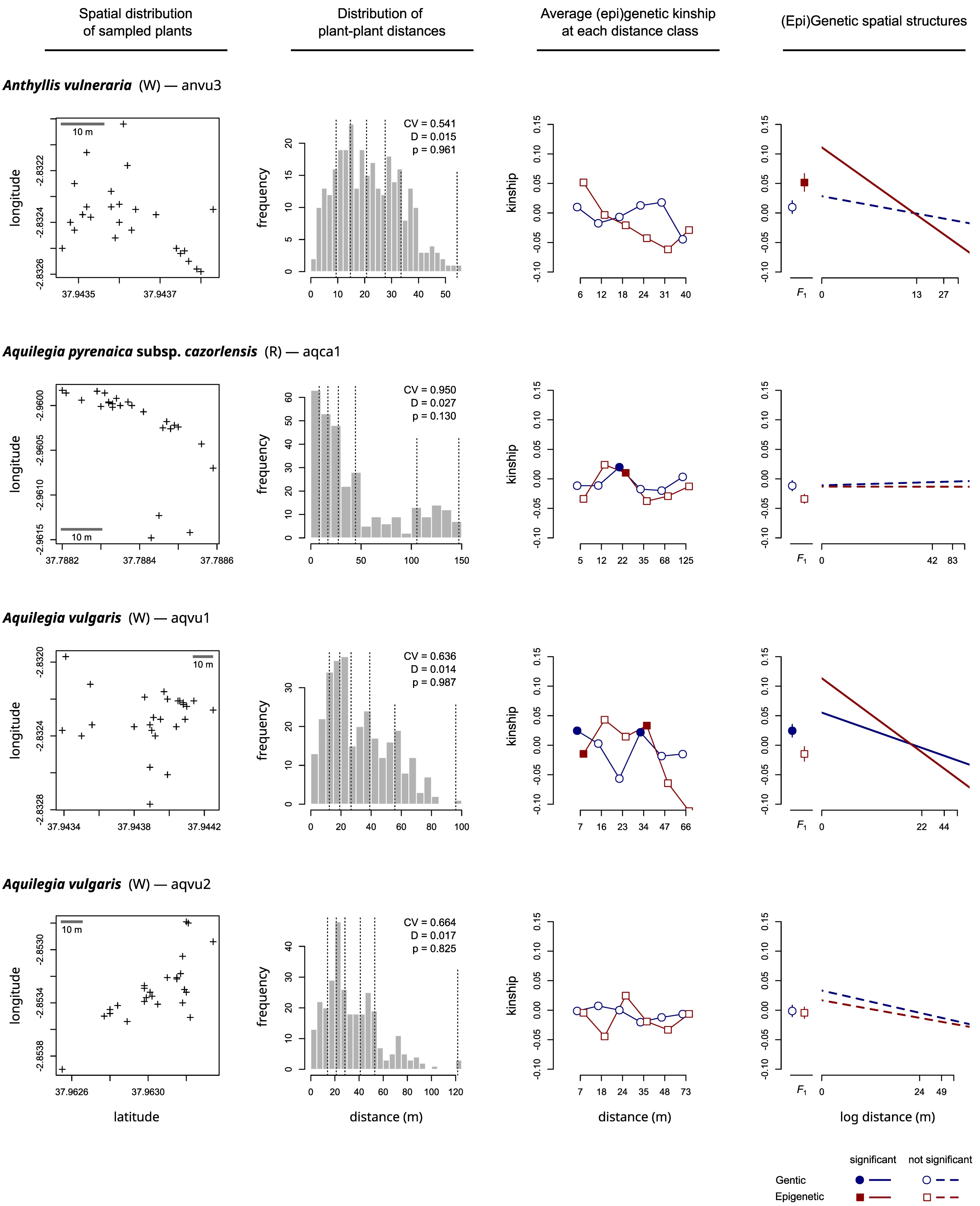

**Figure S3 (3/10).-** Spatial distribution of sampled individuals and spatial genetic and epigenetic structure within study populations.

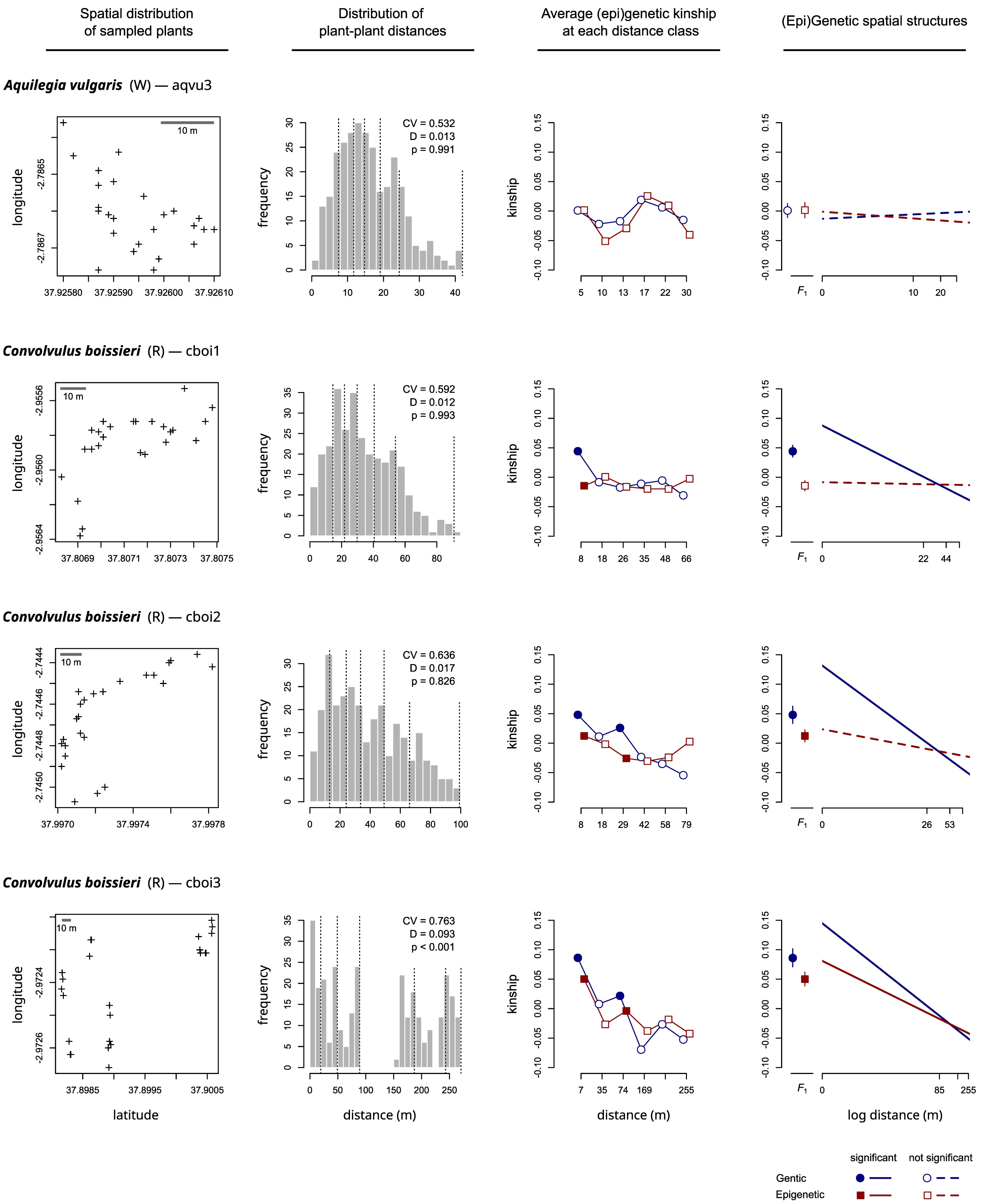

**Figure S3 (4/10).-** Spatial distribution of sampled individuals and spatial genetic and epigenetic structure within study populations.

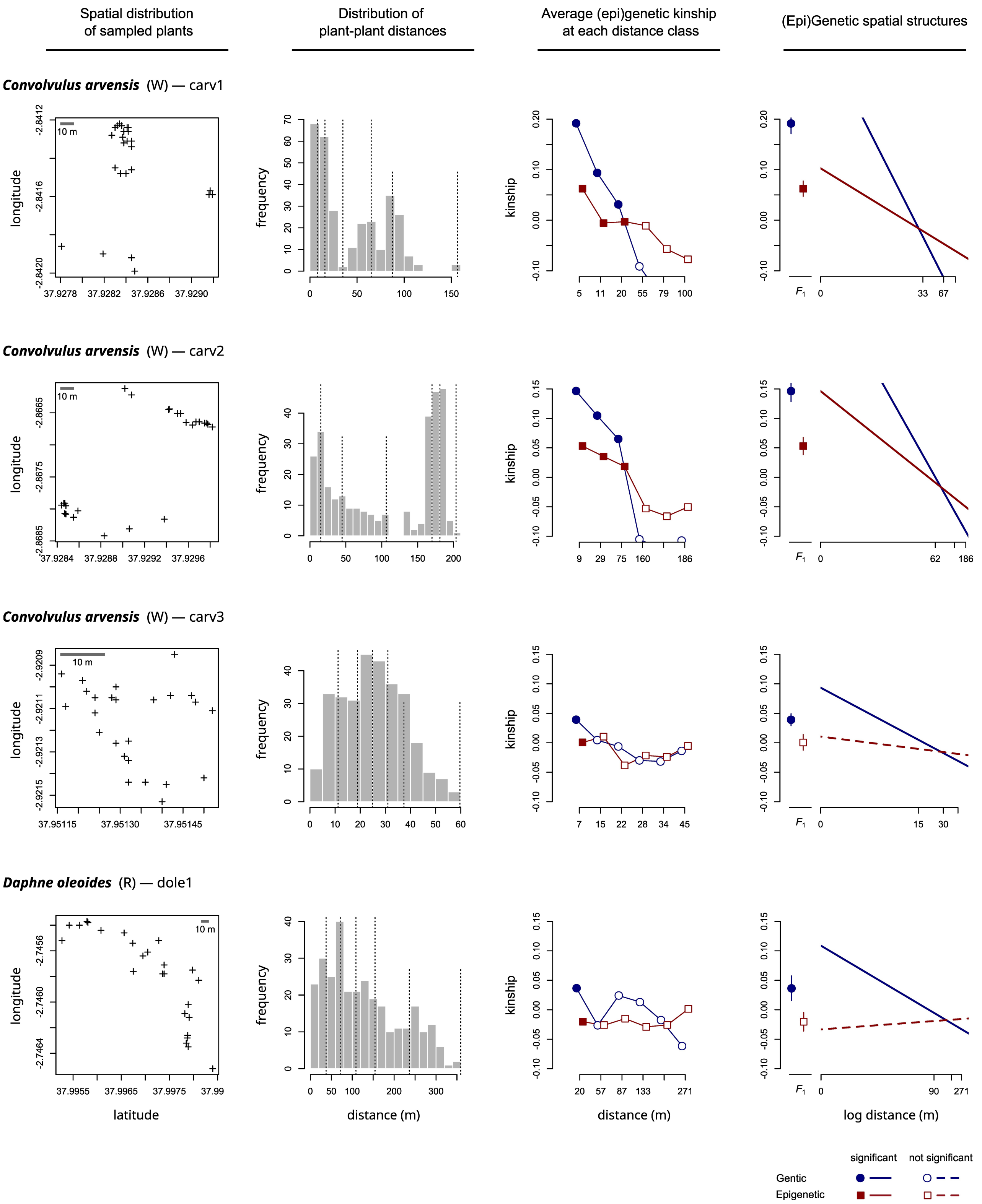

**Figure S3 (5/10).-** Spatial distribution of sampled individuals and spatial genetic and epigenetic structure within study populations.

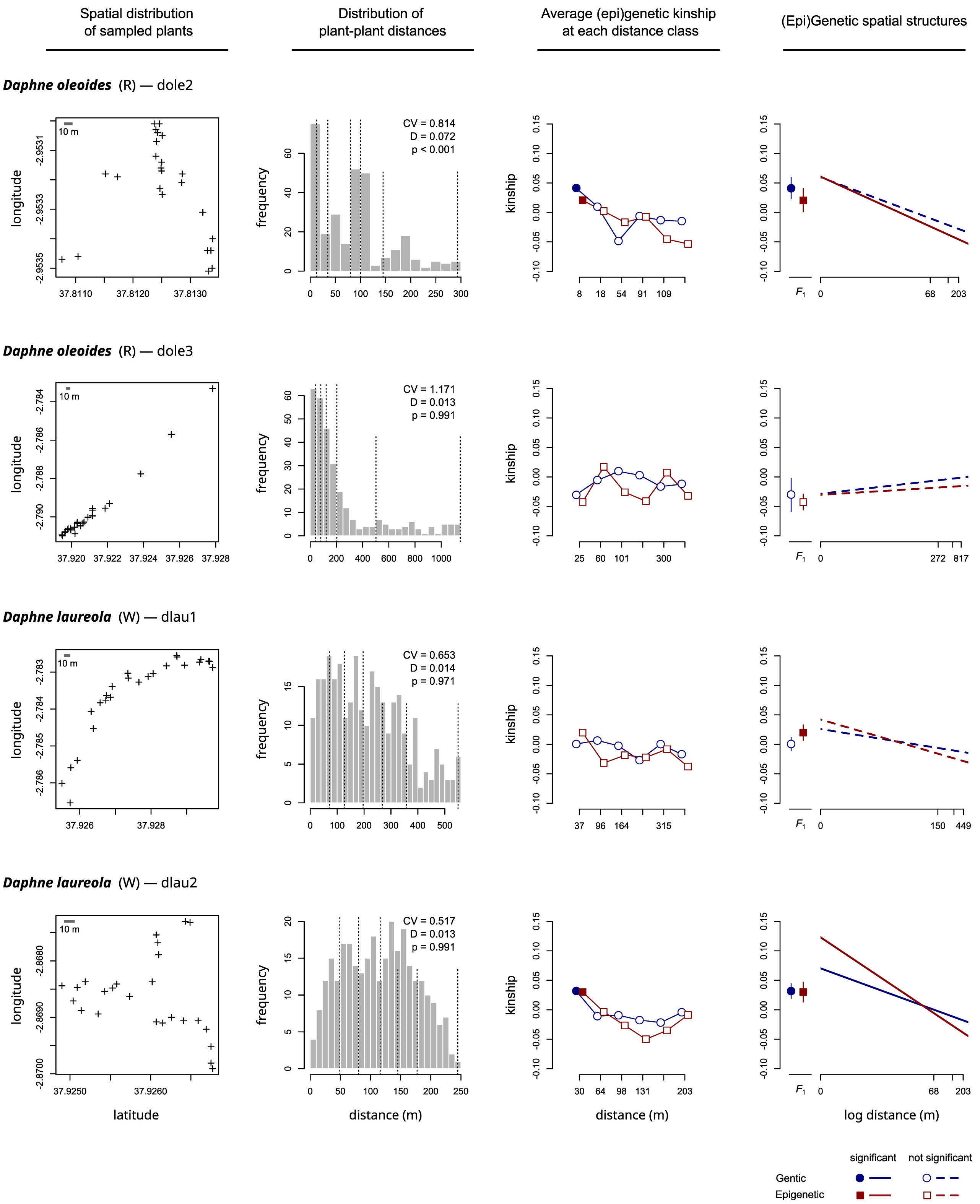

**Figure S3 (6/10).-** Spatial distribution of sampled individuals and spatial genetic and epigenetic structure within study populations.

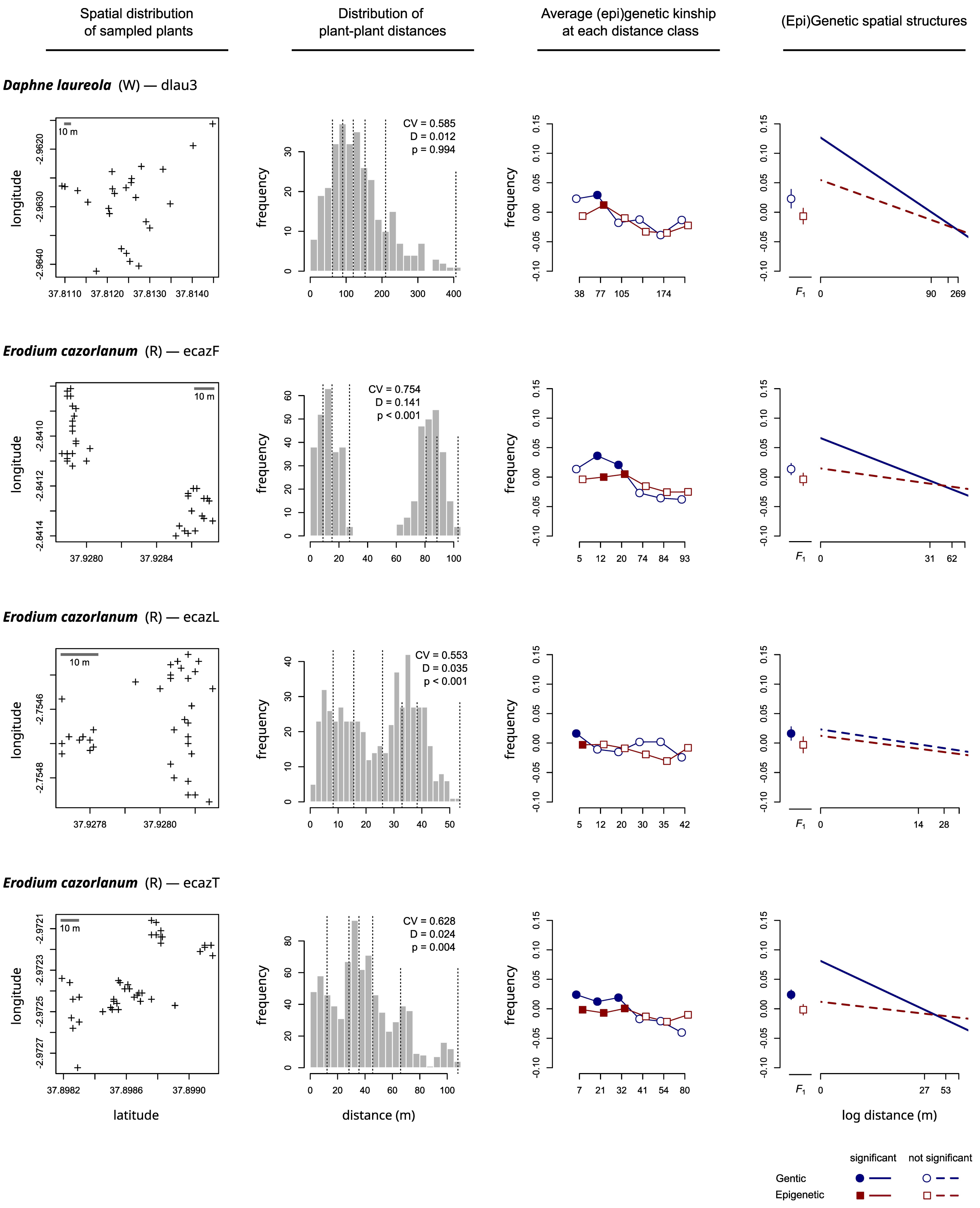

**Figure S3 (7/10).-** Spatial distribution of sampled individuals and spatial genetic and epigenetic structure within study populations.

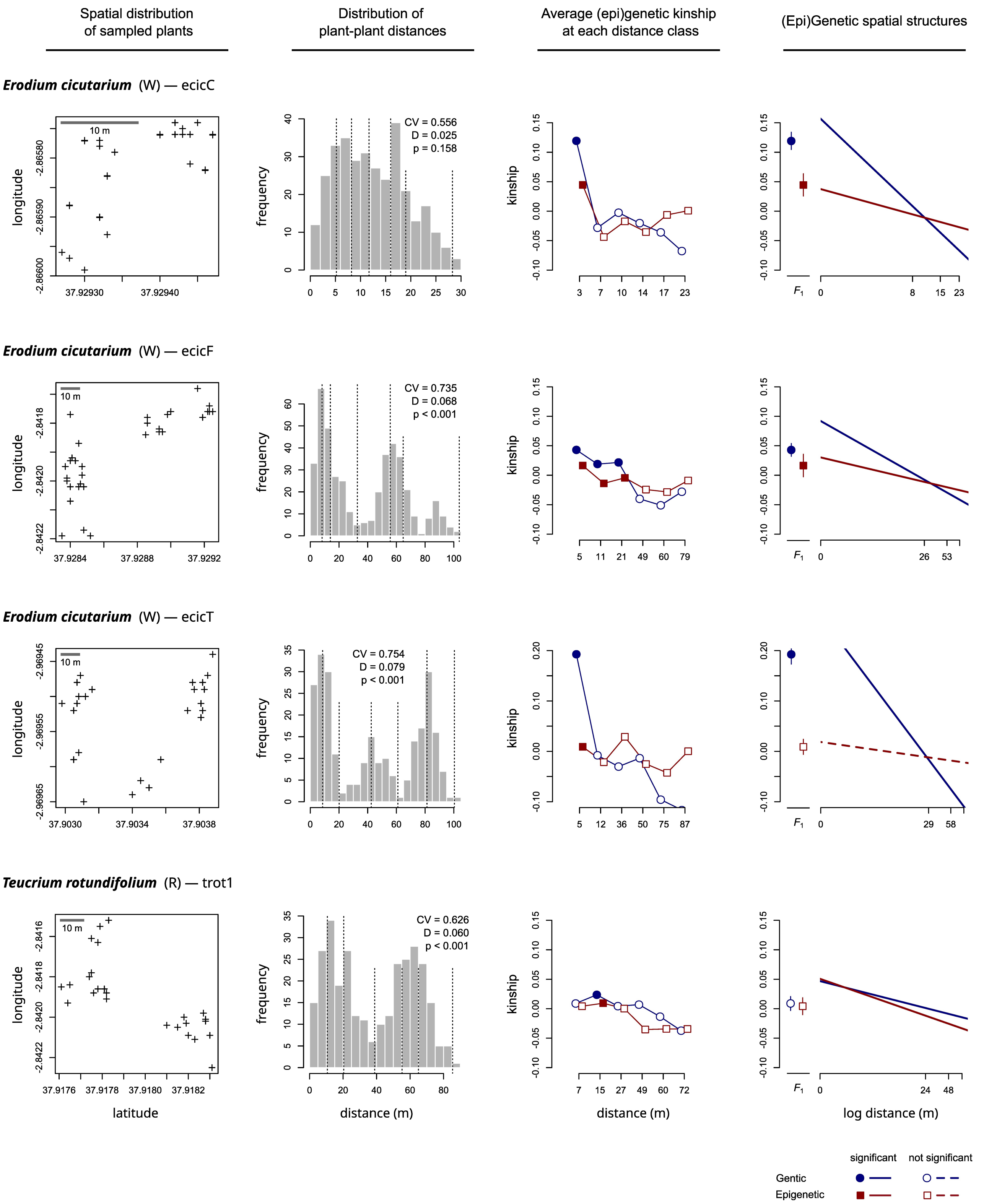

**Figure S3 (8/10).-** Spatial distribution of sampled individuals and spatial genetic and epigenetic structure within study populations.

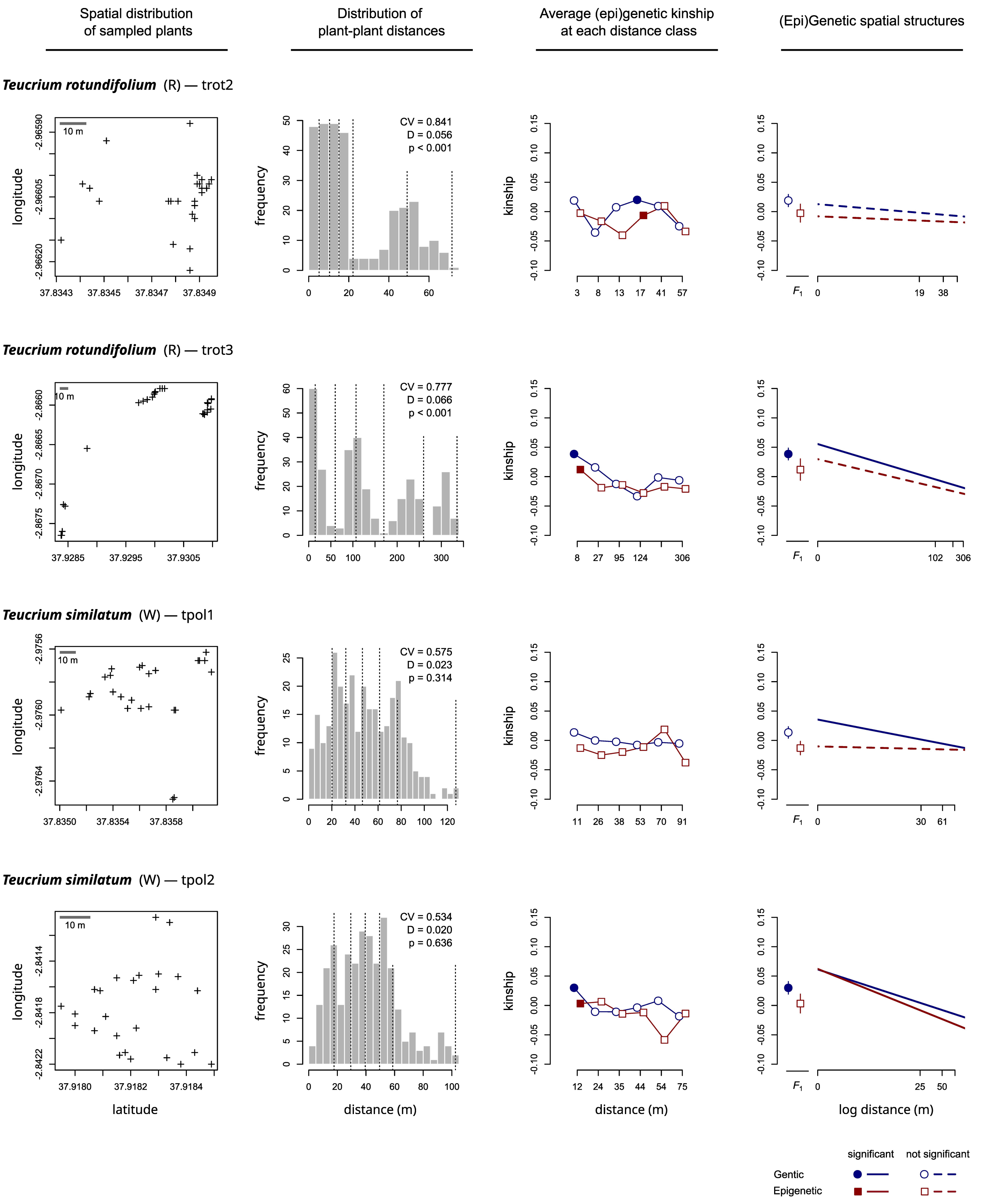

**Figure S3 (9/10).-** Spatial distribution of sampled individuals and spatial genetic and epigenetic structure within study populations.

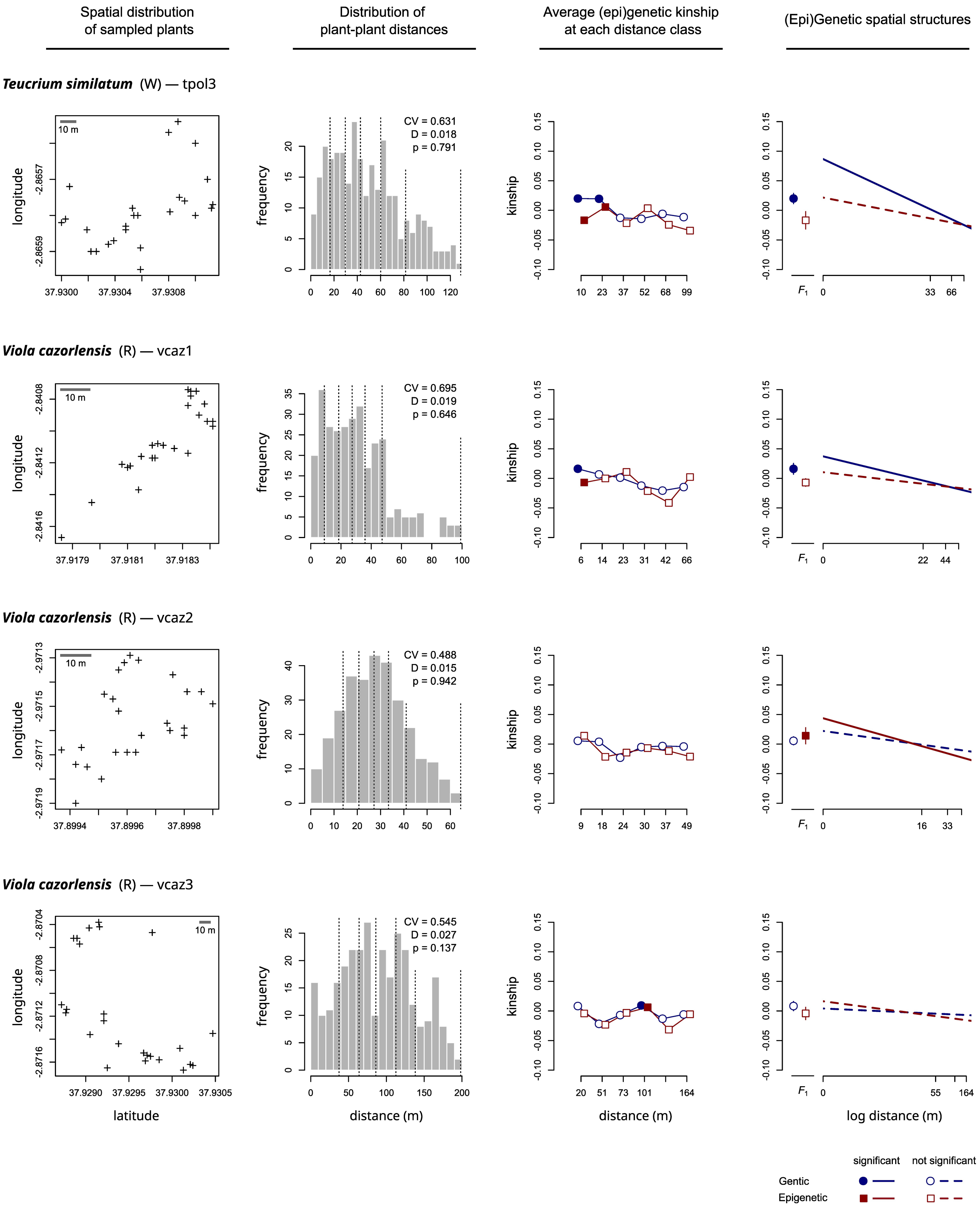

**Figure S3 (10/10).-** Spatial distribution of sampled individuals and spatial genetic and epigenetic structure within study populations.

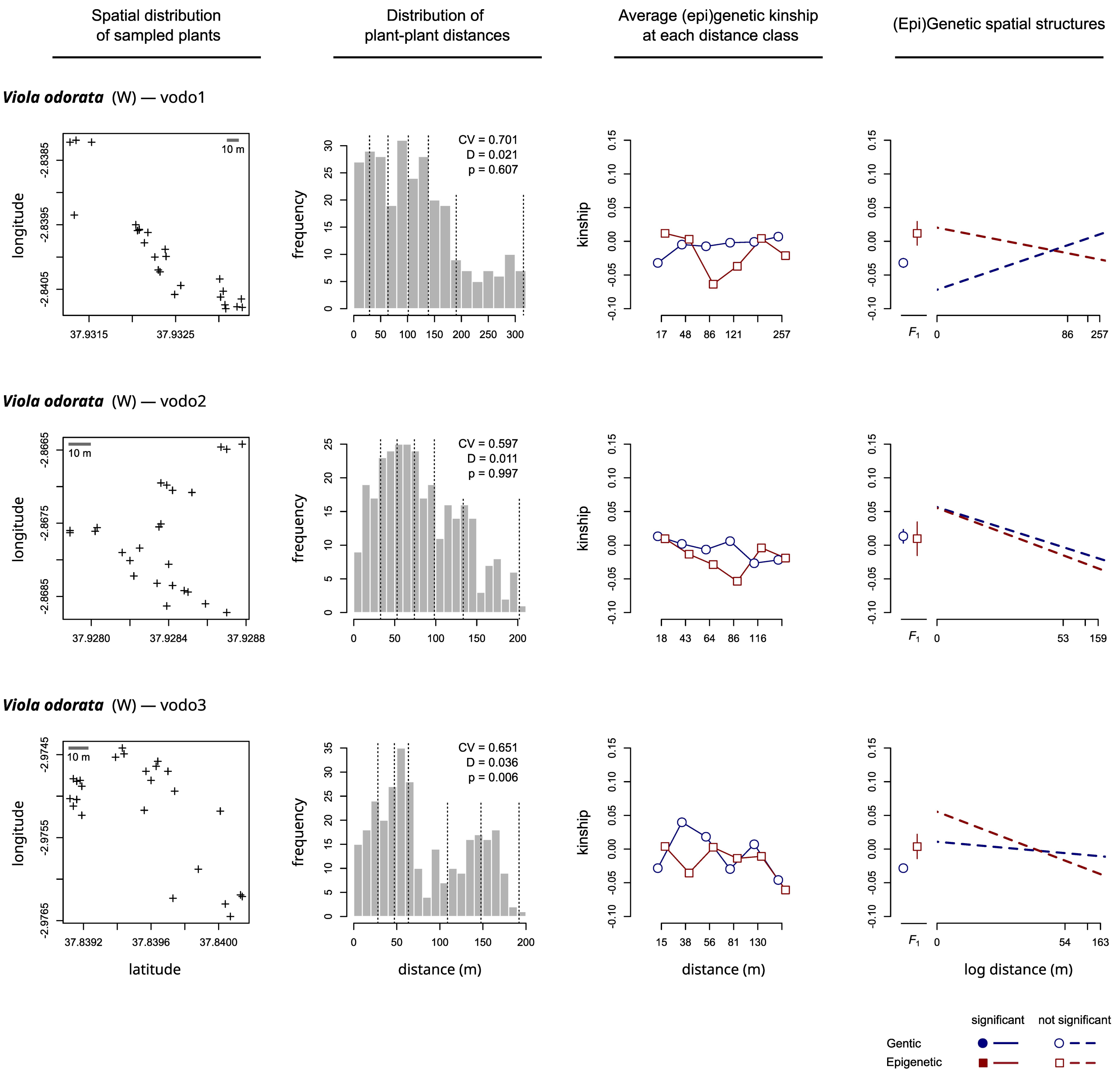
