## Supplementary Tables for "Comparative epigenetic and genetic spatial structure in Mediterranean mountain plants: a multispecific comparison"

**Table S1.-** Distribution and habitat types of the study species.

| Taxa | Distribution type | Habitat type | Distribution |
|---|---|---|---|
| *Anthyllis ramburii* | Narrow endemic | High mountain dwarf procumbent scrubs, sandy dolomitic soils. | Baetic Ranges |
| *Anthyllis vulneraria* | Widespread | Dry grasslands and rocky environments with calcareous soils, broad altitudinal range. | European, Mediterranean basin east to the Caucasus |
| *Aquilegia pyrenaica* subsp. *cazorlensis* | Narrow endemic | Rifts of limestone outcrops, sandy soils in shady, damp sites at cliff bases. | Baetic Ranges |
| *Aquilegia vulgaris* | Widespread | Stream margins, poorly drained open meadows around springs, broad altitudinal range | European temperate element and Mediterranean |
| *Convolvulus boissieri* | Narrow endemic | High mountain dwarf procumbent scrubs, sandy dolomitic soils | Baetic Ranges |
| *Convolvulus arvensis* | Widespread | Cultivated areas, wasteland, roadsides, grassy slopes, broad altitudinal range | Temperate and tropical regions, except Australia |
| *Daphne oleoides* | Narrow endemic | Calcareous xerophytic woodlands and scrublands, rocky slopes | North Africa, southern Europe, Asia Minor |
| *Daphne laureola* | Widespread | Sclerophyllous and semi-deciduous forests, understory of montane forests, mostly in basic soils | Palaearctic |
| *Erodium cazorlanum* | Narrow endemic | High mountain dry grasslands and rockyenvironments, sandy dolomitic soils | Baetic Ranges |
| *Erodium cicutarium* | Widespread | Meadows, flood plains, gravel areas, roadsides and disturbed areas, broad altitudinal range | Eurosiberian Southern-temperate element, circumpolar invasive |
| *Teucrium rotundifolium* | Narrow endemic | Rocky outcrops | Iberian Peninsula–Morocco |
| *Teucrium similatum* | Widespread | Sclerophyllous and semi-deciduous forests, calcareous scrublands and grasslands, rocky slopes, broad altitudinal range | Iberian Peninsula |
| *Viola cazorlensis* | Narrow endemic | Rocky outcrops, cliffs, sandy dolomitic soils | Baetic Ranges |
| *Viola odorata* | Widespread | Open woodlands, hedge banks and scrublands, edges of forests and clearings, broad altitudinal range | Europe and Asia |

**Table S2.-** Primer combinations used for AFLP and MSAP analyses in the seven genera considered in this study.

| | Primer combination | |
|---|---|---|
| Genus | AFLP | MSAP |
| *Anthyllis* | *Mse*I + CGT / *Pst*I + AA | *Mse*I + CAT / *Hpa*II–*Msp*I + TT |
| | *Mse*I + CCT / *Pst*I + AT | *Mse*I + CTC / *Hpa*II–*Msp*I + TA |
| | *Mse*I + CAT / *Pst*I + AC | *Mse*I + CAC / *Hpa*II–*Msp*I + TC |
| | *Mse*I + CAT / *Pst*I + AG | *Mse*I + CCT / *Hpa*II–*Msp*I + TC |
| *Aquilegia* | *Mse*I + CGT / *Pst*I + AA | *Mse*I + CAT / *Hpa*II–*Msp*I + TT |
| | *Mse*I + CAC / *Pst*I + AT | *Mse*I + CTC / *Hpa*II–*Msp*I + TA |
| | *Mse*I + CTT / *Pst*I + AC | *Mse*I + CAC / *Hpa*II–*Msp*I + TC |
| | *Mse*I + CAT / *Pst*I + AG | *Mse*I + CCT / *Hpa*II–*Msp*I + TC |
| *Convolvulus* | *Mse*I + CAC / *Pst*I + AA | *Mse*I + CGC / *Hpa*II–*Msp*I + TA |
| | *Mse*I + CTT / *Pst*I + AT | *Mse*I + CTC / *Hpa*II–*Msp*I + TA |
| | *Mse*I + CTT / *Pst*I + AC | *Mse*I + CGT / *Hpa*II–*Msp*I + TG |
| | *Mse*I + CAT / *Pst*I + AG | *Mse*I + CCT / *Hpa*II–*Msp*I + TC |
| *Daphne* | *Mse*I + CAC / *Pst*I + AA | *Mse*I + CGC / *Hpa*II–*Msp*I + TC |
| | *Mse*I + CAC / *Pst*I + AT | *Mse*I + CTC / *Hpa*II–*Msp*I + TA |
| | *Mse*I + CAT / *Pst*I + AC | *Mse*I + CGA / *Hpa*II–*Msp*I + TG |
| | *Mse*I + CTC / *Pst*I + AG | *Mse*I + CTC / *Hpa*II–*Msp*I + TC |
| *Erodium* | *Mse*I + CGT / *Pst*I + AA | *Mse*I + CAT / *Hpa*II–*Msp*I + TT |
| | *Mse*I + CAC / *Pst*I + AT | *Mse*I + CGC / *Hpa*II–*Msp*I + TA |
| | *Mse*I + CTT / *Pst*I + AC | *Mse*I + CCT / *Hpa*II–*Msp*I + TC |
| | *Mse*I + CTC / *Pst*I + AG | |
| *Teucrium* | *Mse*I + CGC / *Pst*I + AA | *Mse*I + CCT / *Hpa*II–*Msp*I + TT |
| | *Mse*I + CCT / *Pst*I + AT | *Mse*I + CGC / *Hpa*II–*Msp*I + TA |
| | *Mse*I + CTT / *Pst*I + AC | *Mse*I + CAC / *Hpa*II–*Msp*I + TC |
| | *Mse*I + CTT / *Pst*I + AG | |
| *Viola* | *Mse*I + CCC / *Pst*I + AA | *Mse*I + CAC / *Hpa*II–*Msp*I + TC |
| | *Mse*I + CCT / *Pst*I + AT | *Mse*I + CTC / *Hpa*II–*Msp*I + TA |
| | *Mse*I + CTT / *Pst*I + AC | *Mse*I + CGT / *Hpa*II–*Msp*I + TG |
| | *Mse*I + CAT / *Pst*I + AG | *Mse*I + CCT / *Hpa*II–*Msp*I + TC |

**Table S3.-** Population genetic and epigenetic diversity. For each of the populations studied the number of sampled individuals and the estimated inbreeding coefficients (*F*i) are shown. For the genetic (AFLP) and epigenetic (MSAP) markers the number of loci studied, the proportion of these which were polymorphic are denoted. The expected heterozygosity and the associated standard error is also shown for each marker.

| | | | Genetic (AFLP markers) | | | Epigenetic (MSAP markers) | | |
|---|---|---|---|---|---|---|---|---|
| Population | Sampled individuals | *F*i | Num. loci | Proportion polymorphic | Expected heterozygosity | Num. loci | Proportion polymorphic | Expected heterozygosity |
| *Anthyllis ramburii* (Narrow endemic) | | | | | | | | |
| anra1 | 25 | 0.219 | 147 | 0.56 | 0.148 ± 0.013 | 121 | 0.64 | 0.159 ± 0.014 |
| anra2 | 25 | 0.125 | 147 | 0.52 | 0.164 ± 0.014 | 121 | 0.60 | 0.156 ± 0.014 |
| anra3 | 25 | 0.408 | 147 | 0.44 | 0.150 ± 0.014 | 121 | 0.63 | 0.158 ± 0.014 |
| *Anthyllis vulneraria* (Widespread species) | | | | | | | | |
| anvu1 | 25 | 0.402 | 234 | 0.33 | 0.090 ± 0.007 | 204 | 0.65 | 0.189 ± 0.011 |
| anvu2 | 25 | 0.785 | 234 | 0.39 | 0.101 ± 0.008 | 204 | 0.67 | 0.191 ± 0.01 |
| anvu3 | 25 | 0.663 | 234 | 0.37 | 0.109 ± 0.008 | 204 | 0.64 | 0.172 ± 0.009 |
| *Aquilegia pyrenaica* subsp. *cazorlensis* (Narrow endemic) | | | | | | | | |
| aqca1 | 25 | 0.062 | 163 | 0.47 | 0.161 ± 0.015 | 191 | 0.56 | 0.189 ± 0.012 |
| aqca2 | 25 | 0.087 | 163 | 0.49 | 0.146 ± 0.013 | 191 | 0.49 | 0.148 ± 0.012 |
| aqca3 | 25 | 0.113 | 163 | 0.46 | 0.164 ± 0.013 | 191 | 0.65 | 0.218 ± 0.013 |
| *Aquilegia vulgaris* (Widespread species) | | | | | | | | |
| aqvu1 | 25 | 0.149 | 151 | 0.46 | 0.161 ± 0.014 | 152 | 0.68 | 0.225 ± 0.015 |
| aqvu2 | 25 | 0.031 | 151 | 0.53 | 0.165 ± 0.014 | 152 | 0.63 | 0.212 ± 0.015 |
| aqvu3 | 25 | 0.029 | 151 | 0.48 | 0.140 ± 0.013 | 152 | 0.63 | 0.212 ± 0.014 |
| *Convolvulus boissieri* (Narrow endemic) | | | | | | | | |
| cboi1 | 25 | 0.077 | 184 | 0.43 | 0.155 ± 0.013 | 194 | 0.51 | 0.159 ± 0.01 |
| cboi2 | 28 | 0.025 | 184 | 0.40 | 0.128 ± 0.012 | 194 | 0.51 | 0.148 ± 0.011 |
| cboi3 | 25 | 0.032 | 184 | 0.42 | 0.14 ± 0.012 | 194 | 0.39 | 0.142 ± 0.012 |
| *Convolvulus arvensis* (Widespread species) | | | | | | | | |
| carv1 | 25 | 0.121 | 223 | 0.32 | 0.128 ± 0.011 | 157 | 0.31 | 0.127 ± 0.012 |
| carv2 | 25 | 0.090 | 223 | 0.58 | 0.211 ± 0.012 | 157 | 0.54 | 0.153 ± 0.011 |
| carv3 | 25 | 0.162 | 223 | 0.70 | 0.228 ± 0.012 | 157 | 0.47 | 0.141 ± 0.011 |
| *Daphne oleoides* (Narrow endemic) | | | | | | | | |
| dole1 | 25 | 0.124 | 86 | 0.24 | 0.127 ± 0.015 | 95 | 0.54 | 0.214 ± 0.016 |
| dole2 | 25 | 0.044 | 86 | 0.29 | 0.116 ± 0.017 | 95 | 0.56 | 0.198 ± 0.016 |
| dole3 | 25 | 0.130 | 86 | 0.38 | 0.131 ± 0.018 | 95 | 0.50 | 0.191 ± 0.017 |
| *Daphne laureola* (Widespread species) | | | | | | | | |
| dlau1 | 25 | 0.092 | 114 | 0.57 | 0.191 ± 0.017 | 145 | 0.72 | 0.236 ± 0.015 |
| dlau2 | 25 | 0.018 | 114 | 0.52 | 0.137 ± 0.014 | 145 | 0.70 | 0.254 ± 0.014 |
| dlau3 | 25 | 0.015 | 114 | 0.52 | 0.121 ± 0.015 | 145 | 0.65 | 0.213 ± 0.014 |
| *Erodium cazorlanum* (Narrow endemic) | | | | | | | | |
| ecazF | 32 | 0,197 | 162 | 0.63 | 0.227 ± 0.014 | 147 | 0.62 | 0.186 ± 0.011 |
| ecazL | 33 | 0.369 | 162 | 0.53 | 0.176 ± 0.014 | 147 | 0.61 | 0.165 ± 0.011 |
| ecazT | 40 | 0.253 | 162 | 0.65 | 0.198 ± 0.013 | 147 | 0.69 | 0.18 ± 0.011 |
| *Erodium cicutarium* (Widespread species) | | | | | | | | |
| ecicC | 28 | 0.036 | 128 | 0.59 | 0.232 ± 0.015 | 100 | 0.29 | 0.096 ± 0.011 |
| ecicF | 30 | 0.148 | 128 | 0.77 | 0.291 ± 0.016 | 100 | 0.34 | 0.108 ± 0.013 |
| ecicT | 23 | 0.039 | 128 | 0.85 | 0.268 ± 0.015 | 100 | 0.38 | 0.085 ± 0.011 |

**Table S3 (cont.).-** Population genetic and epigenetic diversity. For each of the populations studied the number of sampled individuals and the estimated inbreeding coefficients (*F*i) are shown. For the genetic (AFLP) and epigenetic (MSAP) markers the number of loci studied, the proportion of these which were polymorphic are denoted. The expected heterozygosity and the associated standard error is also shown for each marker.

| Population | Sampled individuals | *F*i | Genetic (AFLP markers) Num. loci | Genetic (AFLP markers) Prop. loci polymorphic | Genetic (AFLP markers) Expected heterozygosity | Epigenetic (MSAP markers) Num. loci | Epigenetic (MSAP markers) Prop. loci polymorphic | Epigenetic (MSAP markers) Expected heterozygosity |
|---|---|---|---|---|---|---|---|---|
| *Teucrium rotundifoium* (Narrow endemic) | | | | | | | | |
| trot1 | 25 | 0.940 | 319 | 0.36 | 0.120 ± 0.008 | 186 | 0.67 | 0.209 ± 0.01 |
| trot2 | 25 | 0.962 | 319 | 0.35 | 0.125 ± 0.008 | 186 | 0.44 | 0.155 ± 0.011 |
| trot3 | 25 | 0.963 | 319 | 0.40 | 0.117 ± 0.007 | 186 | 0.55 | 0.172 ± 0.011 |
| *Teucrium similatum* (Widespread species) | | | | | | | | |
| tpol1 | 25 | 0.965 | 418 | 0.43 | 0.139 ± 0.007 | 213 | 0.49 | 0.132 ± 0.007 |
| tpol2 | 25 | 0.955 | 418 | 0.38 | 0.131 ± 0.007 | 213 | 0.47 | 0.13 ± 0.007 |
| tpol3 | 25 | 0.955 | 418 | 0.45 | 0.143 ± 0.007 | 213 | 0.49 | 0.13 ± 0.007 |
| *Viola cazorlensis* (Narrow endemic) | | | | | | | | |
| vcaz1 | 25 | 0.014 | 261 | 0.41 | 0.12 ± 0.009 | 185 | 0.27 | 0.095 ± 0.008 |
| vcaz2 | 25 | 0.053 | 261 | 0.42 | 0.12 ± 0.008 | 185 | 0.24 | 0.074 ± 0.008 |
| vcaz3 | 25 | 0.053 | 261 | 0.39 | 0.126 ± 0.009 | 185 | 0.21 | 0.07 ± 0.008 |
| *Viola odorata* (Widespread species) | | | | | | | | |
| vodo1 | 24 | 0.306 | 316 | 0.45 | 0.147 ± 0.009 | 107 | 0.52 | 0.157 ± 0.014 |
| vodo2 | 25 | 0.613 | 316 | 0.47 | 0.147 ± 0.009 | 107 | 0.73 | 0.188 ± 0.013 |
| vodo3 | 25 | 0.528 | 316 | 0.67 | 0.185 ± 0.009 | 107 | 0.50 | 0.153 ± 0.015 |

**Table S4.-** Regional-scale genetic and epigenetic structure. For each species the Wright's fixation index ($F_{ST}$) estimated for the genetic markers and for the epigenetic markers are shown. These correspond to estimates using the across-populations averaged inbreeding ($F_i$). Values between brakets denote $F_{ST}$ estimates using inbreeding values of 0 and 1. $F_{ST}$ values in bold indicate significance based on a permutation test.

| Species | Distribution range | $F_i$ | Genetic $F_{ST}$ (AFLP) | Epigenetic $F_{ST}$ (MSAP) |
|---|---|---|---|---|
| *Anthyllis ramburii* | Narrow endemic | 0.32 | **0.059** [**0.044** – **0.078**] | **0.028** [**0.020** – **0.047**] |
| *Anthyllis vulneraria* | Widespread species | 0.62 | 0.008 [0.007 – 0.010] | **0.031** [**0.020** – **0.037**] |
| *Aquilegia pyrenaica* subsp. *cazorlensis* | Narrow endemic | 0.17 | **0.24** [**0.215** – **0.297**] | **0.159** [**0.141** – **0.202**] |
| *Aquilegia vulgaris* | Widespread species | 0.11 | **0.273** [**0.257** – **0.355**] | **0.367** [**0.340** – **0.449**] |
| *Convolvulus boissieri* | Narrow endemic | 0.09 | **0.114** [**0.101** – **0.163**] | **0.056** [**0.044** – **0.073**] |
| *Convolvulus arvensis* | Widespread species | 0.16 | **0.195** [**0.179** – **0.24**] | **0.091** [**0.079** – **0.109**] |
| *Daphne oleoides* | Narrow endemic | 0.19 | **0.022** [0.032 – **0.008**] | 0.018 [0.03 – 0.014] |
| *Daphne laureola* | Widespread species | 0.10 | **0.216** [**0.219** – **0.237**] | **0.081** [**0.078** – **0.123**] |
| *Erodium cazorlanum* | Narrow endemic | 0.44 | **0.139** [**0.105** – **0.161**] | **0.035** [0.027 – **0.049**] |
| *Erodium cicutarium* | Widespread species | 0.21 | **0.207** [**0.188** – **0.226**] | **0.042** [**0.034** – **0.060**] |
| *Teucrium rotundifolium* | Narrow endemic | 0.96 | **0.041** [**0.02** – **0.041**] | **0.030** [**0.013** – **0.031**] |
| *Teucrium similatum* | Widespread species | 0.96 | **0.016** [**0.005** – **0.017**] | **0.008** [0 – **0.009**] |
| *Viola cazorlensis* | Narrow endemic | 0.04 | **0.024** [**0.029** – **0.032**] | **0.012** [**0.010** – **0.019**] |
| *Viola odorata* | Widespread species | 0.62 | **0.023** [0.015 – **0.029**] | **0.021** [0.017 – **0.022**] |

**Table S5.-** Regional-scale genetic and epigenetic structure. For each study species, the results of the analysis of molecular variance (AMOVA) for the genetic (AFLP) and methylation sensitive epigenetic markers (MSAP) are shown. Statistics include, mean squared deviations (*MS*), variance estimates for each level and the associated significance after comparing with random estimates (500 permutations). Note that the degree of freedoms (*df*) are the same for analyses on both markers.

| | | AFLP | | | M-MSAP | | |
|---|---|---|---|---|---|---|---|
| source of variation | *df* | *MS* | variance | *P*-value | *MS* | variance | *P*-value |
| *Anthyllis ramburii* (Narrow endemic) | | | | | | | |
| among populations | 2 | 79.15 | 1.10 | 0.001 | 103.73 | 1.62 | 0.001 |
| individuals within populations | 72 | 870.72 | 12.09 | | 815.68 | 11.33 | |
| *Anthyllis vulneraria* (Widespread species) | | | | | | | |
| among populations | 2 | 32.21 | 0.12 | 0.097 | 144.77 | 2.02 | 0.001 |
| individuals within populations | 72 | 950.32 | 13.20 | | 1572.96 | 21.85 | |
| *Aquilegia pyrenaica* subsp. *cazorlensis* (Narrow endemic) | | | | | | | |
| among populations | 2 | 288.93 | 5.32 | 0.001 | 329.25 | 5.89 | 0.001 |
| individuals within populations | 72 | 833.52 | 11.58 | | 1248.88 | 17.35 | |
| *Aquilegia vulgaris* (Widespread species) | | | | | | | |
| among populations | 2 | 319.01 | 5.98 | 0.001 | 755.84 | 14.51 | 0.001 |
| individuals within populations | 72 | 717.52 | 9.97 | | 1091.92 | 15.17 | |
| *Convolvulus boissieri* (Widespread species) | | | | | | | |
| among populations | 2 | 173.31 | 2.83 | 0.001 | 165.22 | 2.50 | 0.001 |
| individuals within populations | 75 | 986.06 | 13.15 | | 1319.93 | 17.60 | |
| *Convolvulus arvensis* (Widespread species) | | | | | | | |
| among populations | 2 | 381.33 | 6.86 | 0.001 | 176.99 | 2.98 | 0.001 |
| individuals within populations | 72 | 1388.56 | 19.29 | | 1013.28 | 14.07 | |
| *Daphne oleoides* (Narrow endemic) | | | | | | | |
| among populations | 2 | 10.85 | 0.04 | 0.145 | 68.91 | 1.11 | 0.001 |
| individuals within populations | 72 | 318.32 | 4.42 | | 482.32 | 6.70 | |
| *Daphne laureola* (Widespread species) | | | | | | | |
| among populations | 2 | 133.68 | 2.40 | 0.001 | 182.03 | 3.10 | 0.001 |
| individuals within populations | 72 | 495.44 | 6.88 | | 973.12 | 13.52 | |
| *Erodium cazorlanum* (Narrow endemic) | | | | | | | |
| among populations | 2 | 257.23 | 3.23 | 0.001 | 99.52 | 0.97 | 0.001 |
| individuals within populations | 102 | 1642.94 | 16.11 | | 1636.90 | 16.05 | |
| *Erodium cicutarium* (Widespread species) | | | | | | | |
| among populations | 2 | 280.04 | 4.68 | 0.001 | 39.25 | 0.48 | 0.001 |
| individuals within populations | 78 | 1123.39 | 14.40 | | 519.34 | 6.66 | |
| *Teucrium rotundifolium* (Narrow endemic) | | | | | | | |
| among populations | 2 | 87.49 | 0.97 | 0.001 | 112.08 | 1.59 | 0.001 |
| individuals within populations | 72 | 1411.04 | 19.60 | | 1173.04 | 16.29 | |
| *Teucrium similatum* (Widespread species) | | | | | | | |
| among populations | 2 | 80.83 | 0.48 | 0.001 | 87.47 | 1.16 | 0.001 |
| individuals within populations | 72 | 2038.00 | 28.31 | | 1059.60 | 14.72 | |
| *Viola cazorlensis* (Narrow endemic) | | | | | | | |
| among populations | 2 | 50.24 | 0.47 | 0.001 | 83.47 | 1.24 | 0.001 |
| individuals within populations | 72 | 961.28 | 13.35 | | 780.32 | 10.84 | |
| *Viola odorata* (Widespread species) | | | | | | | |
| among populations | 2 | 88.48 | 0.78 | 0.008 | 79.09 | 1.23 | 0.001 |
| individuals within populations | 71 | 1783.52 | 25.12 | | 651.11 | 9.17 | |

**Table S6.-** Fine-scale genetic and epigenetic structure in the study populations. For each population estimates of the slope of the relationship between the logarithm of the spatial distance and the genetic and epigenetic kinships are shown, toghether with the associated statistic and p-value.

| | $log_{10}$(distance)-genetic relatedness | | | $log_{10}$(distance)-epigenetic similarity | | |
|---|---|---|---|---|---|---|
| population | estimate | *t* | *p* | estimate | *t* | *p* |
| *Anthyllis ramburii* (Narrow endemic) | | | | | | |
| anra1 | -0.035 | -4.308 | 0.000 | -0.031 | -2.737 | 0.007 |
| anra2 | -0.010 | -0.901 | 0.369 | 0.000 | -0.011 | 0.991 |
| anra3 | – | – | – | – | – | – |
| *Anthyllis vulneraria* (Widespread species) | | | | | | |
| anvu1 | -0.027 | -3.260 | 0.001 | -0.005 | -0.434 | 0.665 |
| anvu2 | -0.046 | -5.507 | 0.000 | -0.042 | -5.246 | 0.000 |
| anvu3 | -0.011 | -1.090 | 0.277 | -0.045 | -3.340 | 0.001 |
| *Aquilegia pyrenaica* subsp. *cazorlensis* (Narrow endemic) | | | | | | |
| aqca1 | 0.001 | 0.247 | 0.805 | 0.000 | -0.005 | 0.996 |
| aqca2 | – | – | – | – | – | – |
| aqca3 | – | – | – | – | – | – |
| *Aquilegia vulgaris* (Widespread species) | | | | | | |
| aqvu1 | -0.019 | -1.993 | 0.047 | -0.041 | -2.705 | 0.007 |
| aqvu2 | -0.012 | -1.727 | 0.085 | -0.009 | -0.785 | 0.433 |
| aqvu3 | 0.003 | 0.310 | 0.757 | -0.005 | -0.286 | 0.775 |
| *Convolvulus boissieri* (Narrow endemic) | | | | | | |
| cboi1 | -0.028 | -5.075 | 0.000 | -0.001 | -0.172 | 0.864 |
| cboi2 | -0.040 | -7.859 | 0.000 | -0.010 | -1.883 | 0.061 |
| cboi3 | -0.035 | -11.359 | 0.000 | -0.022 | -5.777 | 0.000 |
| *Convolvulus arvensis* (Widespread species) | | | | | | |
| carv1 | -0.114 | -11.273 | 0.000 | -0.035 | -4.915 | 0.000 |
| carv2 | -0.084 | -12.803 | 0.000 | -0.038 | -7.663 | 0.000 |
| carv3 | -0.033 | -3.975 | 0.000 | -0.008 | -0.887 | 0.376 |
| *Daphne oleoides* (Narrow endemic) | | | | | | |
| dole1 | -0.025 | -3.041 | 0.003 | 0.003 | 0.405 | 0.686 |
| dole2 | -0.016 | -1.832 | 0.068 | -0.020 | -3.111 | 0.002 |
| dole3 | 0.004 | 0.343 | 0.732 | 0.002 | 0.303 | 0.762 |
| *Daphne dlaureola* (Widespread species) | | | | | | |
| dlau1 | -0.006 | -0.848 | 0.397 | -0.012 | -1.409 | 0.160 |
| dlau2 | -0.017 | -2.012 | 0.045 | -0.030 | -3.083 | 0.002 |
| dlau3 | -0.028 | -2.599 | 0.010 | -0.015 | -1.439 | 0.151 |
| *Erodium cazorlanum* (Narrow endemic) | | | | | | |
| ecazF | -0.021 | -5.288 | 0.000 | -0.008 | -1.937 | 0.053 |
| ecazL | -0.010 | -1.499 | 0.134 | -0.008 | -1.373 | 0.170 |
| ecazT | -0.025 | -6.538 | 0.000 | -0.006 | -1.516 | 0.130 |
| *Erodium cicutarium* (Widespread species) | | | | | | |
| ecicC | -0.071 | -7.483 | 0.000 | -0.021 | -2.574 | 0.010 |
| ecicF | -0.031 | -4.794 | 0.000 | -0.013 | -2.275 | 0.023 |
| ecicT | -0.085 | -10.428 | 0.000 | -0.009 | -1.593 | 0.113 |

**Table S6 (cont.).-** Fine-scale genetic and epigenetic structure in the study populations. For each population estimates of the slope of the relationship between the logarithm of the spatial distance and the genetic and epigenetic kinships are shown, toghether with the associated statistic and p-value.

| | $log_{10}$(distance)-genetic relatedness | | | $log_{10}$(distance)-epigenetic similarity | | |
|---|---|---|---|---|---|---|
| population | estimate | *t* | *p* | estimate | *t* | *p* |
| *Teucrium rotundifoium* (Narrow endemic) | | | | | | |
| trot1 | -0.014 | -2.646 | 0.009 | -0.020 | -2.493 | 0.013 |
| trot2 | -0.005 | -0.958 | 0.339 | -0.002 | -0.315 | 0.753 |
| trot3 | -0.013 | -3.504 | 0.001 | -0.010 | -1.820 | 0.070 |
| *Teucrium similatum* (Widespread species) | | | | | | |
| tpol1 | -0.010 | -2.078 | 0.039 | -0.001 | -0.156 | 0.876 |
| tpol2 | -0.018 | -2.658 | 0.008 | -0.022 | -2.153 | 0.032 |
| tpol3 | -0.024 | -4.529 | 0.000 | -0.010 | -1.329 | 0.185 |
| *Viola cazorlensis* (Narrow endemic) | | | | | | |
| vcaz1 | -0.013 | -2.711 | 0.007 | -0.006 | -0.975 | 0.330 |
| vcaz2 | -0.008 | -1.276 | 0.203 | -0.017 | -2.133 | 0.034 |
| vcaz3 | -0.002 | -0.507 | 0.612 | -0.006 | -1.034 | 0.302 |
| *Viola odorata* (Widespread species) | | | | | | |
| vodo1 | 0.015 | 1.436 | 0.152 | -0.009 | -0.681 | 0.497 |
| vodo2 | -0.015 | -1.003 | 0.317 | -0.018 | -1.282 | 0.201 |
| vodo3 | -0.004 | -0.230 | 0.818 | -0.018 | -1.103 | 0.271 |

**Table S7.-** Model selection on the effect of the type of distribution on the relationship between the logarithm of the spatial distance and the genetic and epigenetic kinships. Reduced models include the coefficient of variation and the maximum of the spatial distance between sampled plants as explanatory variables. Full mixed models are those in which the type of marker is included as explanatory variable. In each case the best model (in bold) is selected based on the Akaike and Bayes information criteria (AIC and BIC) and significance tests.

| model | AIC | BIC | *p* |
|---|---|---|---|
| Narrow endemics | | | |
| reduced | -213.03 | -205.11 | 0.031 |
| **full (fixed slopes)** | **-215.69** | **-206.19** | **0.167** |
| full (random slopes) * | -215.25 | -202.58 | |
| Widespread species | | | |
| reduced | -190.64 | -181.95 | 0.078 |
| **full (fixed slopes)** | **-191.75** | **-181.32** | **0.004** |
| full (random slopes) * | -198.59 | -184.69 | |

* singular model

**Table S8.-** Direct and indirect effects of the structured equation models for each study species. Values between brackets denote 95% CI calculated after 1000 bootstrap steps. Significant effects are highlighted in bold. Note that these analyses were not performed for ANRA3, AQCA2 and AQCA3 given the lack of spatial location data.

| population | direct effects | | | indirect effects |
|---|---|---|---|---|
| | log(spatial distance) ↓ genetic kinship | log(spatial distance) ↓ epigenetic kinship | genetic kinship ↓ epigenetic kinship | log(spatial distance) ↓ epigenetic kinship |
| *Anthyllis ramburii* (Narrow endemic) | | | | |
| anra1 | **-0.24 [-0.36 – -0.12]** | -0.09 [-0.21 – 0.02] | **0.24 [0.11 – 0.35]** | **-0.06 [-0.12 – -0.02]** |
| anra2 | -0.12 [-0.25 – 0.03] | -0.04 [-0.17 – 0.09] | **0.21 [0.10 – 0.33]** | -0.03 [-0.07 – 0.00] |
| anra3 | – | – | – | – |
| *Anthyllis vulneraria* (Widespread species) | | | | |
| anvu1 | **-0.19 [-0.29 – -0.06]** | 0.02 [-0.10 – 0.14] | **0.23 [0.11 – 0.32]** | **-0.04 [-0.08 – -0.01]** |
| anvu2 | **-0.30 [-0.41 – -0.20]** | **-0.14 [-0.25 – -0.03]** | **0.44 [0.36 – 0.56]** | **-0.14 [-0.19 – -0.09]** |
| anvu3 | -0.06 [-0.19 – 0.07] | **-0.17 [-0.28 – -0.05]** | **0.39 [0.30 – 0.49]** | -0.03 [-0.08 – 0.02] |
| *Aquilegia pyrenaica* subsp. *cazorlensis* (Narrow endemic) | | | | |
| aqca1 | 0.01 [-0.12 – 0.13] | 0.00 [-0.11 – 0.13] | **0.14 [0.04 – 0.26]** | 0.00 [-0.02 – 0.02] |
| aqca2 | – | – | – | – |
| aqca3 | – | – | – | – |
| *Aquilegia vulgaris* (Widespread species) | | | | |
| aqvu1 | **-0.12 [-0.21 – -0.01]** | **-0.11 [-0.20 – -0.01]** | **0.43 [0.33 – 0.51]** | **-0.05 [-0.093 – -0.01]** |
| aqvu2 | -0.04 [-0.18 – 0.13] | -0.02 [-0.13 – 0.10] | **0.36 [0.26 – 0.46]** | -0.01 [-0.07 – 0.04] |
| aqvu3 | 0.02 [-0.09 – 0.13] | -0.03 [-0.12 – 0.05] | **0.59 [0.51 – 0.66]** | 0.01 [-0.06 – 0.08] |
| *Convolvulus boissieri* (Narrow endemic) | | | | |
| cboi1 | **-0.28 [-0.4 – -0.14]** | -0.02 [-0.12 – 0.09] | -0.02 [-0.13 – 0.10] | 0.01 [-0.03 – 0.04] |
| cboi2 | **-0.41 [-0.5 – -0.31]** | -0.08 [-0.18 – 0.02] | 0.04 [-0.06 – 0.14] | -0.02 [-0.06 – 0.03] |
| cboi3 | **-0.55 [-0.64 – -0.44]** | **-0.25 [-0.37 – -0.14]** | 0.02 [-0.09 – 0.14] | -0.01 [-0.08 – 0.05] |
| *Convolvulus arvensis* (Widespread species) | | | | |
| carv1 | **-0.55 [-0.64 – -0.42]** | -0.01 [-0.12 – 0.10] | **0.41 [0.32 – 0.50]** | **-0.22 [-0.28 – -0.15]** |
| carv2 | **-0.60 [-0.66 – -0.51]** | 0.02 [-0.06 – 0.10] | **0.59 [0.50 – 0.67]** | **-0.35 [-0.40 – -0.30]** |
| carv3 | **-0.22 [-0.33 – -0.13]** | 0.00 [-0.11 – 0.11] | **0.23 [0.13 – 0.35]** | **-0.05 [-0.09 – -0.02]** |
| *Daphne oleoides* (Narrow endemic) | | | | |
| dole1 | **-0.17 [-0.29 – -0.06]** | 0.03 [-0.11 – 0.17] | 0.05 [-0.07 – 0.17] | -0.01 [-0.04 – 0.01] |
| dole2 | -0.11 [-0.22 – 0.01] | **-0.14 [-0.26 – -0.04]** | **0.30 [0.19 – 0.41]** | -0.03 [-0.08 – 0.00] |
| dole3 | 0.02 [-0.08 – 0.12] | 0.01 [-0.11 – 0.15] | **0.31 [0.22 – 0.42]** | 0.01 [-0.03 – 0.04] |
| *Daphne laureola* (Widespread species) | | | | |
| dlau1 | -0.05 [-0.15 – 0.07] | -0.06 [-0.17 – 0.05] | **0.43 [0.33 – 0.52]** | -0.02 [-0.06 – 0.03] |
| dlau2 | **-0.12 [-0.24 – -0.01]** | **-0.15 [-0.27 – -0.03]** | **0.24 [0.11 – 0.35]** | **-0.03 [-0.07 – -0.01]** |
| dlau3 | **-0.15 [-0.26 – -0.03]** | -0.03 [-0.15 – 0.09] | **0.34 [0.25 – 0.43]** | **-0.05 [-0.10 – -0.01]** |
| *Erodium cazorlanum* (Narrow endemic) | | | | |
| ecazF | **-0.24 [-0.32 – -0.16]** | -0.04 [-0.12 – 0.05] | **0.19 [0.09 – 0.28]** | **-0.05 [-0.08 – -0.02]** |
| ecazL | -0.07 [-0.15 – 0.03] | -0.06 [-0.13 – 0.03] | 0.02 [-0.05 – 0.10] | 0.00 [-0.01 – 0.00] |
| ecazT | **-0.23 [-0.30 – -0.17]** | -0.03 [-0.11 – 0.04] | **0.09 [0.02 – 0.17]** | **-0.02 [-0.04 – -0.01]** |
| *Erodium cicutarium* (Widespread species) | | | | |
| ecicC | **-0.40 [-0.50 – -0.28]** | -0.03 [-0.14 – 0.08] | **0.29 [0.18 – 0.4]** | **-0.11 [-0.18 – -0.07]** |
| ecicF | **-0.23 [-0.32 – -0.13]** | -0.03 [-0.14 – 0.05] | **0.33 [0.24 – 0.42]** | **-0.07 [-0.12 – -0.04]** |
| ecicT | **-0.55 [-0.65 – -0.45]** | 0.04 [-0.09 – 0.14] | **0.23 [0.11 – 0.36]** | **-0.13 [-0.20 – -0.06]** |

**Table S8 (cont.).-** Direct and indirect effects of the structured equation models for each study species. Values between brackets denote 95% CI calculated after 1000 bootstrap steps. Significant effects are highlighted in bold. Note that these analyses were not performed for ANRA3, AQCA2 and AQCA3 given the lack of spatial location data.

| | direct effects | | | indirect effects |
|---|---|---|---|---|
| population | log(spatial distance) ↓ genetic kinship | log(spatial distance) ↓ epigenetic kinship | genetic kinship ↓ epigenetic kinship | log(spatial distance) ↓ epigenetic kinship |
| *Teucrium rotundifoium* (Narrow endemic) | | | | |
| trot1 | **-0.15 [-0.29 – -0.03]** | **-0.14 [-0.28 – -0.01]** | 0.04 [-0.08 – 0.16] | -0.01 [-0.03 – 0.01] |
| trot2 | -0.06 [-0.19 – 0.07] | -0.02 [-0.14 – 0.09] | 0.06 [-0.05 – 0.16] | 0.00 [-0.03 – 0.00] |
| trot3 | **-0.20 [-0.31 – -0.08]** | -0.08 [-0.23 – 0.05] | 0.10 [-0.01 – 0.21] | **-0.02 [-0.05 – 0.00]** |
| *Teucrium similatum* (Widespread species) | | | | |
| tpol1 | -0.12 [-0.25 – 0.03] | 0.01 [-0.13 – 0.14] | 0.13 [0.00 – 0.24] | -0.02 [-0.05 – 0.00] |
| tpol2 | **-0.15 [-0.27 – -0.04]** | **-0.11 [-0.24 – -0.01]** | 0.06 [-0.07 – 0.19] | -0.01 [-0.04 – 0.01] |
| tpol3 | **-0.25 [-0.48 – -0.10]** | -0.03 [-0.14 – 0.08] | **0.16 [0.05 – 0.28]** | **-0.04 [-0.11 – -0.01]** |
| *Viola cazorlensis* (Narrow endemic) | | | | |
| vcaz1 | **-0.16 [-0.26 – -0.05]** | -0.05 [-0.16 – 0.09] | 0.06 [-0.06 – 0.19] | -0.01 [-0.04 – 0.01] |
| vcaz2 | -0.07 [-0.20 – 0.07] | **-0.12 [-0.26 – 0.00]** | 0.00 [-0.11 – 0.11] | 0.00 [-0.01 – 0.01] |
| vcaz3 | -0.03 [-0.14 – 0.09] | -0.06 [-0.19 – 0.06] | 0.06 [-0.07 – 0.17] | 0.00 [-0.02 – 0.00] |
| *Viola odorata* (Widespread species) | | | | |
| vodo1 | 0.09 [-0.02 – 0.21] | -0.07 [-0.18 – 0.05] | **0.37 [0.24 – 0.51]** | 0.03 [-0.01 – 0.09] |
| vodo2 | -0.06 [-0.18 – 0.07] | -0.07 [-0.22 – 0.05] | 0.06 [-0.06 – 0.22] | 0.00 [-0.02 – 0.00] |
| vodo3 | -0.01 [-0.11 – 0.10] | -0.05 [-0.16 – 0.03] | **0.72 [0.51 – 0.89]** | -0.01 [-0.07 – 0.01] |

**Table S9.-** Model selection on the comparison between narrow endemics and widespread species in each of the causal relationships studied. Reduced models include the coeficient of variation and the maximum of the spatial distance between sampled plants as explanatory variables. Full mixed models add the type of marker as explanatory variable. In each case the best model (in bold) is selected based on the Akaike and Bayes information criteria (AIC and BIC) and significance tests.

| model | AIC | BIC | *p* |
|---|---|---|---|
| *spatial distance → genetic relatedness* | | | |
| **reduced** | **-26.67** | **-18.35** | **0.087** |
| full (fixed slopes) | -22.72 | -12.74 | |
| *genetic relatedness → epigenetic similarity* | | | |
| reduced | -2.69 | 5.62 | <0.001 |
| **full (fixed slopes)** | **-20.01** | **-10.03** | **0.469** |
| full (random slopes) * | -17.44 | -4.13 | |
| *spatial distance → epigenetic similarity* (direct effect) | | | |
| **reduced** | **-98.37** | **-91.71** | **0.029** |
| full | -97.63 | -89.32 | |
| *spatial distance → epigenetic relatedness* (indirect effect) | | | |
| reduced | -74.20 | -65.88 | 0.001 |
| full (fixed slopes) | -76.44 | -66.46 | 0.006 |
| **full (random slopes)** | **-81.68** | **-68.37** | |

* singular model
